# Molecular basis of ubiquitin-independent recognition and degradation of ODC/Antizyme by the 26S proteasome

**DOI:** 10.64898/2026.07.19.739413

**Authors:** Diego R. Ramos-Ortiz, Hao-Hsuan Hsieh, Ken C. Dong, Andreas Martin

**Affiliations:** Biophysics Graduate Group, University of California, Berkeley, CA 94720, USA; California Institute for Quantitative Biosciences, University of California at Berkeley, Berkeley, CA 94720, USA; Howard Hughes Medical Institute, University of California at Berkeley, Berkeley CA, 94720; Department of Molecular and Cell Biology, University of California at Berkeley, Berkeley, CA 94720, USA

## Abstract

The ubiquitin-proteasome system represents the main pathway for targeted protein degradation in eukaryotic cells. The majority of substrates is recruited for degradation through ubiquitin modifications, and the underlying principles are well established. However, the requirements for ubiquitin-independent substrates are still poorly understood. Here, we reveal the mechanisms for the antizyme-mediated degradation of the yeast ornithine decarboxylase (yODC), the first reported ubiquitin-independent substrate of the 26S proteasome. Using biochemical studies and cryo-EM structure determination, we show how antizyme binding makes the yODC monomer prone for degradation by exposing an interface that is normally buried in the catalytically active ODC dimer. Together with a surface on antizyme, yODC forms a two-part interface that binds the N-terminal coiled coil of two ATPase subunits, Rpt4 and Rpt5, for delivery to the 26S proteasome motor. This positions the N-terminal unstructured region of yODC for insertion into the ATPase channel to initiate degradation, which we found does not depend on a specific sequence. Interestingly, binding of the globular yODC/antizyme complex to the Rpt4/Rpt5 coiled coil allosterically stabilizes a proteasome conformation that facilitates substrate engagement by the ATPase motor and may represent a primed pre-initiation state with a general role in ubiquitin-dependent and -independent degradation.

## Introduction

The 26S proteasome is responsible for the targeted degradation of a majority of eukaryotic proteins in the context of general proteostasis, quality control, and the regulatory of numerous cellular processes^1^. Although most proteasome substrates are marked for degradation by poly-ubiquitin chains, there are several examples for ubiquitin-independent degradation ^2–4^, with ornithine decarboxylase (ODC) being the first described more than 30 years ago^5–8^. ODC functions as a homodimeric enzyme and catalyzes the first step in the polyamine biosynthetic pathway, which is essential to maintain normal cell growth^9,10^. This is regulated through an interesting feedback mechanism, where high polyamine concentrations induce the expression of ODC Antizyme (referred to as Antizyme), which promotes the separation of the ODC dimer, formation of an ODC/Antizyme complex, and the ubiquitin-independent degradation of the Antizyme-bound ODC monomer^9–11^. Yeast ODC (yODC) contains an N-terminal unstructured region, whereas mammalian ODC bears a C-terminal tail. Previous work had identified yODC’s N-terminal tail as essential for targeting to the 26S proteasome, and it was suggested that appending this tail to other folded substrates would be sufficient for their degradation in a ubiquitin- and Antizyme-independent manner^12^. However, the molecular basis for this process, in particular the recruitment of ODC to the proteasome, remained largely unknown.

The 26S proteasome is comprised of a barrel-shaped 20S core particle (CP), which contains proteolytic sites in an internal chamber, and a 19S regulatory particle (RP), which mechanically processes protein substrates and transfers them into CP^13^. The RP can be further subdivided into the base and lid subcomplexes. The base includes six distinct ATPase subunits in the clockwise order Rpt1, Rpt2, Rpt6, Rpt3, Rpt4, and Rpt5 that from the heterohexameric AAA+ (ATPases Associated with various cellular Activities) motor of the proteasome, as well as three ubiquitin receptors, Rpn1, Rpn13 and Rpn10^14–19^. The lid with its 9 nine subunits binds to the side and top of the base, and contains the deubiquitinase Rpn11, positioned above the entrance to the ATPase motor^20,21^. To undergo efficient degradation, substrates must have a bipartite degradation signal comprised of (1) a proteasome-binding element, usually in the form of a poly-ubiquitin chain, and (2) an unstructured initiation region of at least 25 amino acids ^22–24^. This flexible region or “tail” is required to reach into the motor for engagement by conserved pore-1 loops of the ATPase subunits, which initiates subsequent translocation, unfolding, and substrate transfer into CP for proteolysis^22,23,25^. During this process, Rpn11 is tasked with cleaving off ubiquitin modifications from the substrates in a co-translocational manner^26–28^. Importantly, more recent characterizations of various ubiquitin-independent systems revealed that those substrates also rely on a bipartite signal for degradation, consisting of a proteasome-binding element and an unstructured region for initiation^3,4,8,29,30^.

Cryo-EM and Cryo-ET studies of the 26S proteasome have revealed an intricate landscape of various conformations that coordinate efficient substrate selection and turnover^31–36^. To initiate degradation, the proteasome adopts a resting or engagement-competent (also called s1) state, in which the entrance to the ATPase motor is well accessible, yet the central channel through the RP is narrow and not coaxially aligned with CP^37,38^. In this conformation, the hexameric motor assumes a fixed spiral-staircase arrangement with Rpt3 at the top, Rpt2 at the bottom, and Rpt6 at an intermediate seam position. Inter-subunit interactions involving the pore-1 loops of four ATPase subunits hold this resting-state staircase, which also seems to prevent ATP hydrolysis^39^. In this resting state, a receptor-bound ubiquitinated substrate can insert its unstructured initiation region into the accessible central channel, where engagement by the pore-1 loops triggers a conformational transition from the resting to a series of ATP-hydrolyzing processing (or non-s1) states. These states are characterized by a ∼30°-rotation of the upper RP, formation of a continuous processing channel, and a repositioning of Rpn11 to partially obstruct the motor entrance for efficient substrate deubiquitination^24,40^. The ATPase subunits in these non-s1 states adopt various spiral staircase arrangements, likely representing snapshots of hand-over-hand movements that are driven by ATP hydrolysis, mechanically unfold substrate proteins, and translocate the unstructured polypeptides into CP^41–43^. While the human 26S proteasome appears most of the time to statically reside in the resting state, the yeast proteasome spontaneously samples the processing states, which leads to a considerable basal ATPase activity^39,40,44^.

Here, we reconstitute the ubiquitin-independent, Antizyme-mediated degradation of yODC by the 26S proteasome and examine the role of yODC’s N-terminal region in this process. Through biochemical assays we show that an unstructured tail is required, yet not sufficient for yODC degradation. In agreement with the model of a bipartite degradation signal, the tail’s sequence is irrelevant, and recruitment for degradation depends on the globular domains of the yODC/Antizyme complex. Using a combination of cryo-EM, AlphaFold3 modeling, and site-directed mutagenesis, we map the yODC/Antizyme binding site to the coiled coil of Rpt4 and Rpt5, a region previously associated with ubiquitin-chain binding^36,45–48^ and allosteric regulation of the proteasome’s conformational state^44^. Remarkably, yODC/Antizyme binding to this coiled coil stabilizes a pre-initiation state in which key s1-specific interactions resolve and the ATPase motor becomes more dynamic, with its pore-1 loops primed for substrate engagement and the transition to non-s1 substrate-processing states.

## Results

### yODC degradation is ubiquitin-independent and requires formation of a stable Antizyme complex

To establish the minimum requirements for the degradation of yODC by the 26S proteasome, we reconstituted this reaction *in vitro*. We used purified endogenous 26S proteasome from *Saccharomyces cerevisiae* and recombinantly-expressed yODC and yeast Antizyme (Az) in *Escherichia coli*. To guarantee our yODC construct maintained its native flexible N-terminus during the purification process, we used an N-terminal fusion with Smt3 that was subsequently cleaved by Ulp1. Purified components were incubated *in vitro*, and the proteasomal degradation was assessed by SDS PAGE (Fig. 1a). In agreement with previous reports, both the yODC homodimer and Antizyme were stable in the presence of 26S proteasome^6,7,12^. Interestingly, pre-incubating yODC with Antizyme did not render it susceptible to degradation, and size-exclusion chromatography (SEC) confirmed that mixing purified yODC and Antizyme was not sufficient to produce a heterodimeric complex (Extended Data Fig. 1a). This suggests that other processes beyond Antizyme expression regulate yODC-dimer disruption and yODC/Antizyme complex formation or that the complex forms co-translationally. Indeed, co-expressing yODC and Antizyme in *E. coli* produced stable yODC/Antizyme heterodimers that allowed yODC degradation *in vitro* (Fig. 1a). Degradation of the entire yODC protein was confirmed by isolating the peptides for LC-MS (Extended Data Fig. 7), showing that 96 % of the yODC-derived products were between 7 and 13 amino acids in length, with a median of 9 amino acids, which is in good agreement with previous studies of proteasome-generated peptide products ^49^.

**Figure 1.**
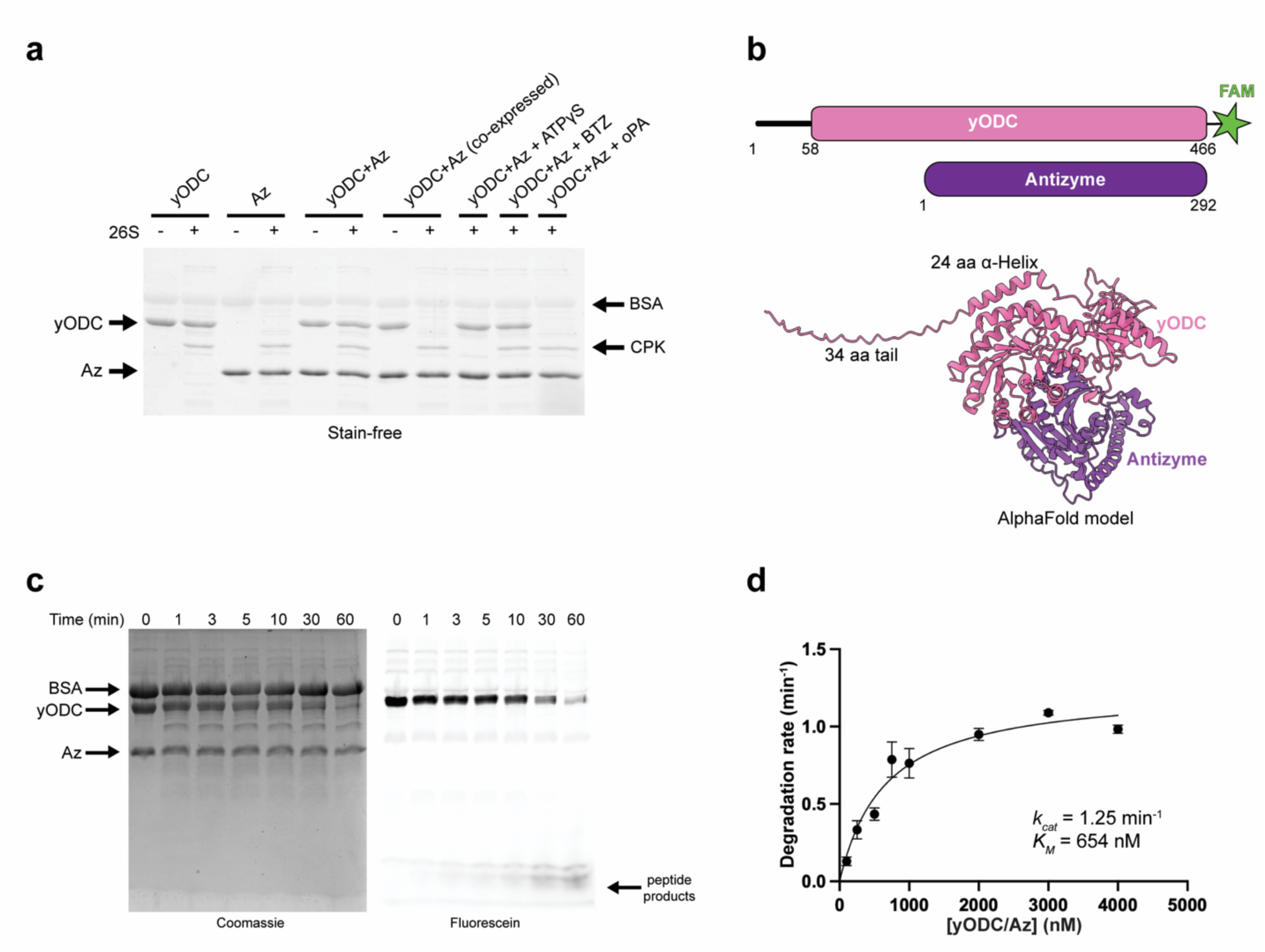
yODC degradation is ubiquitin-independent and requires formation of a stable complex with Antizyme. **a**, SDS-PAGE analysis of yODC degradation by the 26S proteasome. 60-minute end-point assay showing only co-expressed yODC/Antizyme complexes lead to proteasome-mediated degradation of yODC. Control reactions were performed in the presence of the slowly hydrolysable ATP analogue, ATPγS, the proteasome inhibitor bortezomib (BTZ), or the Rpn11 deubiquitination inhibitor o-phenanthroline (oPA). BSA, Bovine serum albumin; CPK, Creatine phosphokinase. **b**, Top, Schematic representation of yODC (pink) and Antizyme (purple). The attachment-point of the FAM (5-Fluoresceinamidite) dye at yODC’s C-terminus is represented by a green star. Bottom, AlphaFold3 prediction of yODC/Antizyme. **c**, SDS-PAGE analysis of samples taken at indicated times from the degradation reaction of FAM-labeled yODC. **d**, Michaelis-Menten analysis of yODC/Antizyme degradation by the 26S proteasome. Initial degradation rates were determined by tracking FAM fluorescence polarization.

Control experiments in the presence of the slowly hydrolyzable ATP analog ATPγS or the proteasome inhibitor bortezomib confirmed that yODC degradation relies on ATP-dependent mechanical unfolding and translocation by the 19S RP, and on the proteolytic cleavage by the 20S CP (Fig. 1a). In contrast, inhibition of the deubiquitinase Rpn11 with o-phenanthroline (oPA) did not interfere with yODC degradation, consistent with this being a ubiquitin-independent process (Fig. 1a).

To increase the throughput and sensitivity of our degradation assay, we sought to identify a suitable position on yODC for attachment of a fluorophore. Previous reports have suggested that yODC’s ubiquitin-independent degron is located at the N-terminus, which is proposed to be unstructured^6,12^. In agreement, AlphaFold3-modeling^42^ of yODC/Antizyme predicted an unstructured stretch of 34 amino acids at yODC’s N-terminus, which led us to attach a Fluorescein-labeled peptide via a sortase reaction to yODC’s C-terminus (Fig. 1b and Extended Data Fig. 1b-d). The degradation of dye-labeled yODC and formation of peptide products was first monitored through time-resolved SDS-PAGE, which indicated complete and processive degradation (Fig. 1c). For

Michaelis-Menten analyses, we measured the changes in fluorescence polarization (FP) of the yODC-attached fluorophore during steady-state degradation at various yODC/Antizyme concentrations, which revealed a substrate affinity of *K_M_* = 654 nM (95% CI [480, 887]) and a degradation rate of *k_cat_* = 1.25 min^-1^ (95% CI [1.13, 1.38]) (Fig. 1d and Extended Data Fig. 1e). This rate is in nice agreement with the degradation velocity of 1.31 min^-1^ determined from the gel-based assay (Fig. 1c). Using these parameters, we designed a series of experiments to examine the basis for Antizyme-dependent yODC degradation at a near-saturating (4 μM) substrate concentration, allowing us to characterize different yODC mutants regarding their degradation rate while remaining sensitive to affinity defects.

### yODC’s unstructured initiation region is sequence-independent and not sufficient for degradation

It has been widely reported that yODC’s N-terminal region (amino acids 1-50) is required for ubiquitin-independent degradation, and appending this sequence to folded proteins destabilizes them *in vivo* ^6,12^. This has led to the conclusion that the native sequence of yODC’s N-terminal region acts as an independent degron and is sufficient to target substrates to the 26S proteasome for degradation. As there are currently no experimentally determined structures of the yODC/Antizyme complex available, we used an AlphaFold3 model as guidance to generate a series of mutations directed at an unstructured stretch of 34 residues (1-34) and a downstream 24-residue α-helix (35-58) in order to dissect the role of yODC’s N-terminal region for degradation (Fig. 1b and 2a). Using the gel-based degradation assay with unlabeled yODC (Fig. 2b and Extended Data Fig. 2a-b), we determined a degradation rate of ∼ 2 min^-1^ for the wild-type protein. This is slightly faster than the rate measured for fluorescently labeled yODC, which may result from the large hydrophobic dye slowing translocation or product release from the CP barrel. We observed that blocking yODC’s free N-terminus with a folded domain, Smt3, or deleting the first 34 amino acids (Δ34yODC) completely abolished degradation, which confirms the requirement for an unstructured initiation region. To test the sequence-dependence of this region, we appended an HA tag (HA), which was proposed to inhibit degradation^6^, or substituted the first 34 amino acids with an *Arbacia punctulata* Cyclin B-derived sequence (cycB34), previously characterized as a suitable unstructured initiation region in the context of ubiquitin-dependent degradation^24^. Remarkably, there was no effect of the HA tag fusion (*k_cat_* = 2.3 min^-1^) and only a modest reduction in rate by the cycB34 replacement (*k_cat_* = 1.4 min^-1^). To rule out that the downstream 24-residue α-helix plays a role in proteasome binding or directing the insertion of the 34-residue tail into the ATPase motor, we substituted this α-helix with a Cyclin B-derived sequence (35cycB58) or made two proline substitutions (2xPro) to disrupt its secondary structure. Degradation rates of 2.5 min^-1^ for 35cycB58 and 2.0 min^-1^ for 2xPro indicated that the N-terminal alpha-helix has no sequence- or structure-specific role in yODC degradation. Taken together, our results confirm that yODC’s N-terminal segment is a sequence-independent, non-specific initiation region for proteasomal degradation. This was further tested in competition assays, where we analyzed the degradation of dye-labeled wild-type yODC in the presence of unlabeled variants with previously described N-terminal mutations (Fig. 2c and Extended Data Fig. 2h). All mutants, except for the truncation Δ34yODC, were able to inhibit degradation to the same extent as unlabeled wild-type yODC, which confirms that the identity of the N-terminal flexible region is largely irrelevant for degradation. Δ34yODC is a weak inhibitor as it lacks an initiation region and cannot be committed for degradation. Smt3-yODC, on the other hand, acted as a competitive inhibitor, despite being degradation-incompetent, likely because the flexible region between Smt3 and the globular domain of yODC can partially enter the central channel as a hairpin, leading to a long enough residence time of this variant to interfere with wild-type yODC degradation.

**Figure 2.**
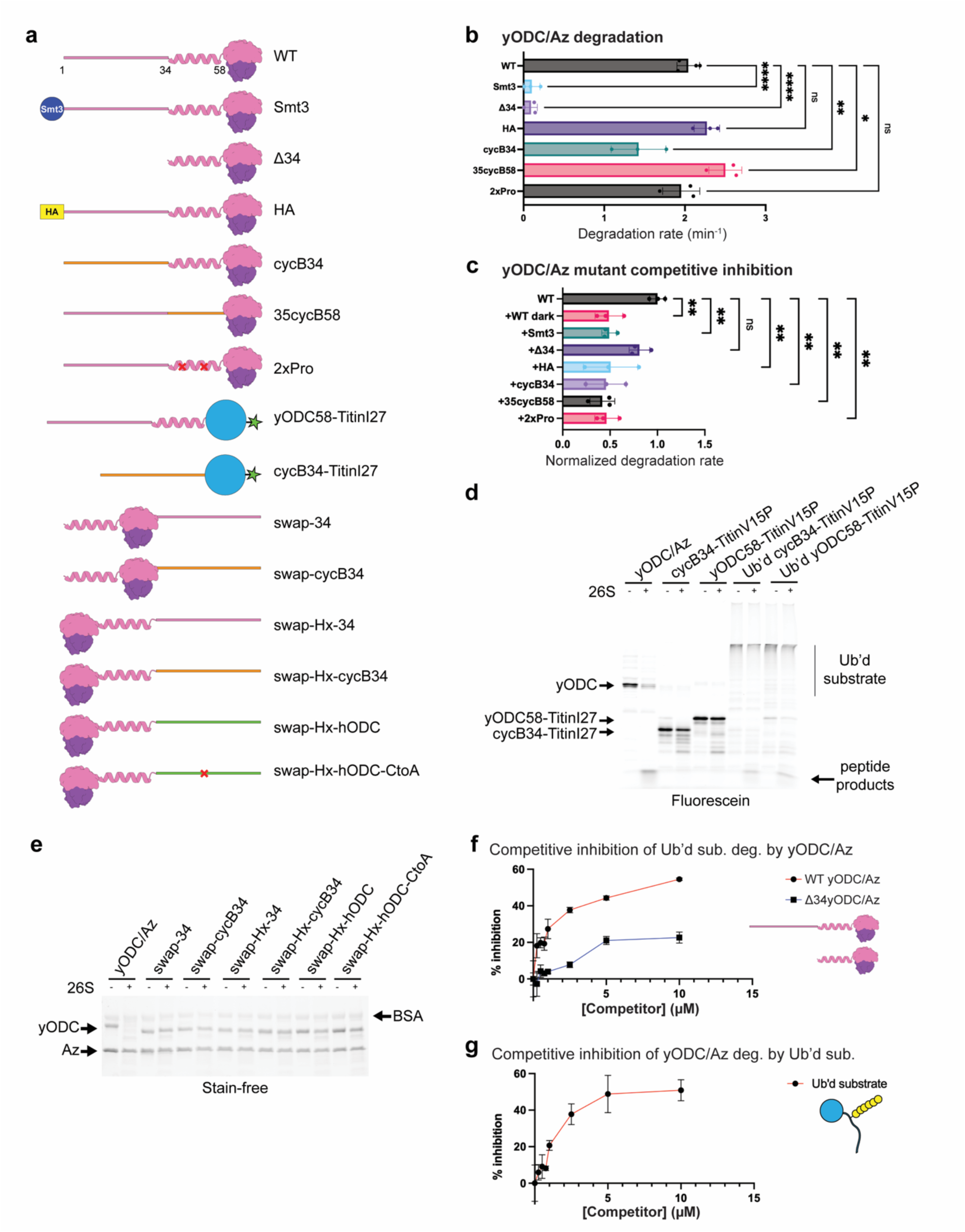
yODC’s unstructured N-terminus is a sequence-independent initiation region and not sufficient for degradation. **a**, Schematic representation of yODC (pink) and Antizyme (purple) complexes illustrating the assayed mutations and modification. Cyclin B- and hODC-derived sequences are shown in orange and green, respectively. Positions of single-amino acid substitutions are marked by a red “X”, and FAM dyes are represented by a green star **b**, Near-saturating multiple-turnover degradation rates for yODC/Antizyme mutants determined by SDS-PAGE-analyzed time courses. Statistical significance was calculated using an ordinary one-way ANOVA with Dunnett’s multiple comparisons test against wild-type yODC/Az (n = 3 technical replicates): ∗∗∗∗p < 0.0001 (Smt3), ∗∗∗∗p < 0.0001 (Δ34), ∗∗p = 0.0042 (cycB34), ∗p = 0.0348 (35cycB58), p = 0.9805 (2xPro), and p = 0.4832 (HA). **c**, Competitive inhibition of FAM-labeled WT yODC/Antizyme degradation by yODC/Antizyme mutants. Initial degradation rates were determined by tracking FAM-fluorescence polarization and normalized to the uninhibited degradation rate. Statistical significance was calculated using an ordinary one-way ANOVA with Dunnett’s multiple comparisons test against wild-type yODC/Az (n = 3 technical replicates): ∗∗p = 0.0069 (+WT dark), ∗∗p = 0.0072 (+Smt3), p = 0.5630 (+Δ34), ∗∗p = 0.0043 (+cycB34), ∗∗p < 0.0024 (+35cycB58), ∗∗p = 0.0047 (+2xPro), and ∗∗p = 0.0093 (+HA). **d**, SDS-PAGE analysis of the 60-min endpoints for the incubations of Cyclin B tail-Titin^V15P^ (cycB34-TitinV15P) or yODC tail-Titin^V15P^ (yODC58-TitinV15P) chimeras with 26S proteasome show that ODC’s unstructured initiation region is not sufficient to target substrates for proteasome-mediated degradation. **e**, SDS-PAGE analysis of yODC tail-location mutants incubated with 26S proteasome. The 30-minute endpoint shows that transplanting the unstructured initiation region to yODC’s C-terminus inhibits proteasome-mediated degradation. **f**, Competitive inhibition of the degradation of FAM-labeled ubiquitinated substrate by WT and Δ34yODC/Antizyme. **g**, Competitive inhibition of the degradation of FAM-labeled yODC/Antizyme by ubiquitinated substrate. Initial degradation rates for FAM-labeled substrates at various competitive inhibitor concentrations (0-10 μM) were determined by tracking the FAM fluorescence polarization and the % inhibition was calculated relative to the uninhibited degradation rate.

Our observations cast doubt on the prevailing model that the N-terminal region alone is sufficient to target yODC or other fused proteins directly to the proteasome for degradation. To address this unambiguously, we built two dye-labeled titin-I27 model substrates with destabilizing V15P mutation (titin^V15P^): yODC58-titin^V15P^, containing yODC’s first 58 residues at the N-terminus, and cycB34-titin^V15P^, containing a N-terminal 34-residue cyclin B-derived initiation region (Fig. 2a and Extended Data Fig. 2c-d). Neither of those substrates got turned over in our gel-based assay (Fig. 2d), but proteasome-dependent degradation could be rescued by ubiquitinating them prior to their incubation with the 26S proteasome (Fig. 2d and Extended Data Fig. 2e-f). This indicates that yODC’s native N-terminus cannot independently target substrates for degradation without another 26S proteasome-binding element. We propose that yODC’s N-terminal tail is not a ubiquitin-independent degron, but instead functions as a flexible initiation region and hence just one component of a bipartite degradation signal, similar to unstructured regions in substrates for ubiquitin-dependent degradation.

### Degradation of yODC cannot initiate from an unstructured region at the C-terminus

We next sought to test whether the position of the initiation region has an effect on yODC degradation. Since the N- and C-terminal residues of the globular yODC portion lie in reasonable proximity, we relocated the initiation region to the C-terminus and characterized the proteasomal turnover in our gel-based assay (Fig. 2a,e). Surprisingly, transplanting yODC’s N-terminal unstructured stretch of 34 amino acids (1-34) to the C-terminus (swap-34) abolished degradation. To rule out sequence-dependent effects, we replaced this C-terminal tail with a 34-residue cyclin B-derived sequence (swap-cycB34), which also prevented degradation (Fig. 2a,e). We wondered whether the 24-residue α-helix (35-58) that follows the native N-terminal tail in yODC is required as a spacer to allow the tail to reach the proteasome motor entrance, and we therefore constructed two more substrates with an insertion of this α-helix between yODC’s original C-terminus and the transplanted 34-residue initiation sequences (swap-Hx-34 and swap-Hx-cycB34). Neither of them got degraded (Fig. 2a,e), even though the distance between the transplantation spots, position 58 and the C-terminal D466, is only 14 A#.

In search for a C-terminal sequence that could rescue yODC degradation, we tested the unstructured C-terminal 37 residues (425-461) of *Homo sapiens* ODC (hODC). Like yODC, hODC was identified as a ubiquitin-independent substrate, but intriguingly, its degron was mapped to the C-terminus ^50^ and is apparently a specific feature of vertebrate ODC (Extended Data Fig. 2g and Extended Data Fig. 6). We therefore constructed a yODC variant with its 24-amino acid α-helix transplanted to the C-terminus followed by the 37-residue tail of hODC (swap-Hx-hODC; Fig. 2a). In addition, we created the same variant with a C to A substitution at the position equivalent to yODC residue 441 (swap-Hx-hODC-CtoA; Fig. 2a), as C441A was previously shown to impair the ubiquitin-independent degradation of hODC^50,51^. Remarkably, neither swap-Hx-hODC nor swap-Hx-hODC-CtoA were degraded by the 26S proteasome (Fig. 2e).

In summary, our results indicate that the 26S proteasome is unable to initiate degradation of yODC from a flexible C-terminal tail. Although it cannot be ruled out that mechanical unfolding of ODC is differentially challenging from the two termini, it is more likely that strict geometric requirements and the orientation of the flexible initiation region determine ODC’s engagement by the ATPase motor. This suggests that there is a dedicated yODC/Antizyme-binding site on the 26S proteasome that orients yODC’s N-terminus for ideal initiation, which is again consistent with the model for bipartite degradation signals.

### yODC/Antizyme competes with ubiquitinated substrates for proteasome binding and degradation

To get initial insight into potential binding sites of yODC/Antizyme on the 26S proteasome, we performed competition experiments with our well-established ubiquitinated titin^V15P^-23-K-35 model substrate. In this substrate, the titin I27 domain with destabilizing V15P mutation is followed by 23 unstructured residues, a single lysine for ubiquitination, and a 35 residue C-terminal tail^24^. We used this substrate in its dye-labeled form to compete with unlabeled yODC or vice versa. Previous studies indicated that hODC competes with ubiquitinated substrates by utilizing or overlapping with ubiquitin-binding sites on the proteasome ^52^. Indeed, when measuring multiple-turnover degradation of the dye-labeled ubiquitinated substrate, we also observed a potent inhibition in the presence of unlabeled wild-type yODC/Antizyme, while the effect was much weaker for the tailless Δ34yODC/Antizyme variant that is unable to engage (Fig. 2f and Extended Data Fig. 2i-j). Similarly, the unlabeled ubiquitinated substrate inhibited the degradation of dye-labeled yODC/Antizyme with comparable potency (Fig. 2g and Extended Data Fig. 2k). These findings suggest that yODC/Antizyme and the ubiquitinated substrate bind to the 26S proteasome with similar affinities and potentially using at least partially overlapping sites.

### yODC/Antizyme is targeted for degradation by binding the Rpt4/Rpt5 coiled coil

After establishing that yODC’s N-terminus serves as an unstructured initiation region and does not directly target yODC for degradation, we reasoned that its globular domain and/or Antizyme must have a dedicated binding site on the proteasome. To identify this site, we utilized cryogenic electron microscopy (cryo-EM) studies of the yODC/Antizyme-bound proteasome. Samples were prepared by manually mixing the 26S proteasome with yODC/Antizyme on ice before plunge freezing after 45 s, assuming that the incubation at low temperature would allow initial binding, but suppress yODC degradation (Extended Data Fig. 3e-f). From the collected dataset, we identified the proteasome in two non-processing conformations: the resting (s1) state and a “primed” (s1’) state that overall resembled the E_B_ conformation previously observed for substrate-engaged h26S proteasome^36^ (Fig. 3a, Extended Data Fig. 8, Table 1). While the s1 state in our sample showed no signs of bound yODC/Antizyme, the s1’ map contained unassigned density on the N-terminal coiled coil formed by two ATPase subunits Rpt4 and Rpt5 (Fig. 3b left). Focused refinement with a mask around this additional density and the Rpt4/Rpt5 coiled coil revealed a low-resolution volume that well accommodates an AlphaFold3 model of yODC/Antizyme bound to the coiled coil (yODC/Antizyme:Rpt4/5cc) (Fig. 3b right, 3c and Extended Data Fig. 3g-i,k).

**Figure 3.**
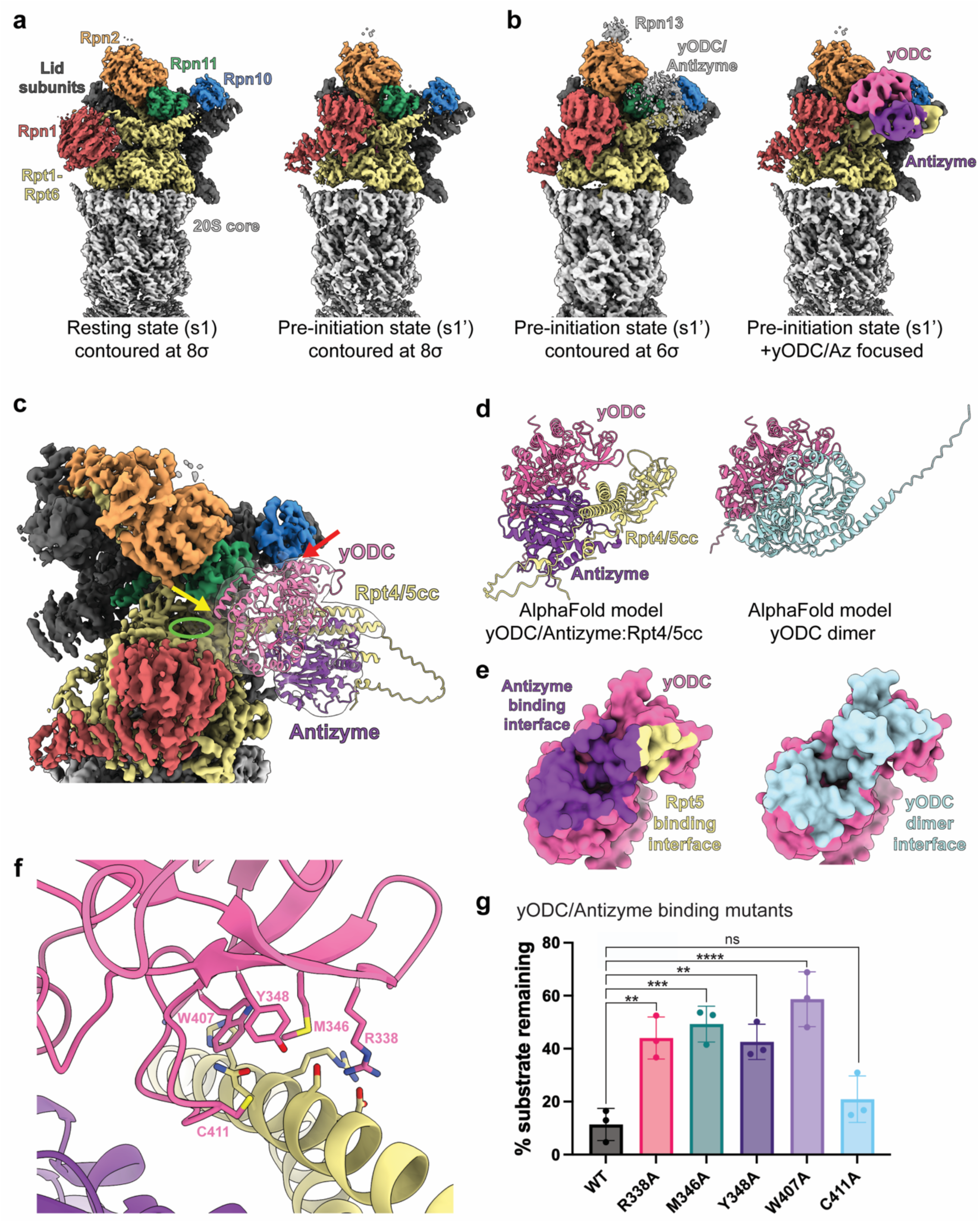
Formation of yODC/Antizyme exposes an interface that binds to the Rpt4/Rpt5 coiled coil. **a**, Cryo-EM density for the resting state (s1, left) and pre-initiation state (s1’, right) proteasomes, with the 20S core in gray and the 19S RP colored by subunits. **b**, Left, cryo-EM map of the s1’ proteasome at low contour reveals unassigned density near the Rpt4/Rpt5 coiled coil. Right, focused cryo-EM map of the additional density corresponding to yODC/Antizyme (pink and purple) overlayed onto the density for the s1’ proteasome. **c**, Cryo-EM density of the s1’ proteasome overlayed with the AlphaFold3 model for yODC/Antizyme bound to Rpt4/Rpt5 coiled coil. The last amino acid of yODC’s N-terminal α-helix at position 35 (yellow arrow) is right above the entrance to the proteasome’s processing channel (green ring). The red arrow indicates yODC’s C-terminus. **d**, Side-by-side view of AlphaFold3 models for the yODC/Antizyme complex bound to the Rpt4/Rpt5 coiled coil (left) and the yODC homodimer (right) reveals overlapping interfaces on yODC, which explains how the Antizyme-mediated yODC degradation depends on separation of the dimer. **e**, Surface representation of yODC (pink) from AlphaFold3 models shown in **d**. Buried surface area is colored according to nearest predicted binding partner using the same color scheme as in **d**. **f**, AlphaFold3-predicted interface between ODC and the Rpt5 N-terminal helix, with the potentially critical residues shown in stick representation. **g**, Substrate remaining after 7.5-min incubation with the 26S proteasome for yODC binding mutants (1 μM), as quantified by SDS-PAGE. Statistical significance was calculated using an ordinary one-way ANOVA with Dunnett’s multiple comparisons test against wild-type yODC/Az (n = 3 technical replicates): ∗∗p = 0.0012 (R338A), ∗∗∗p = 0.0003 (M346A), ∗∗p = 0.0017 (Y348A), ∗∗∗∗p < 0.0001 (W407A), and p = 0.4702 (C411A).

**Table 1.**
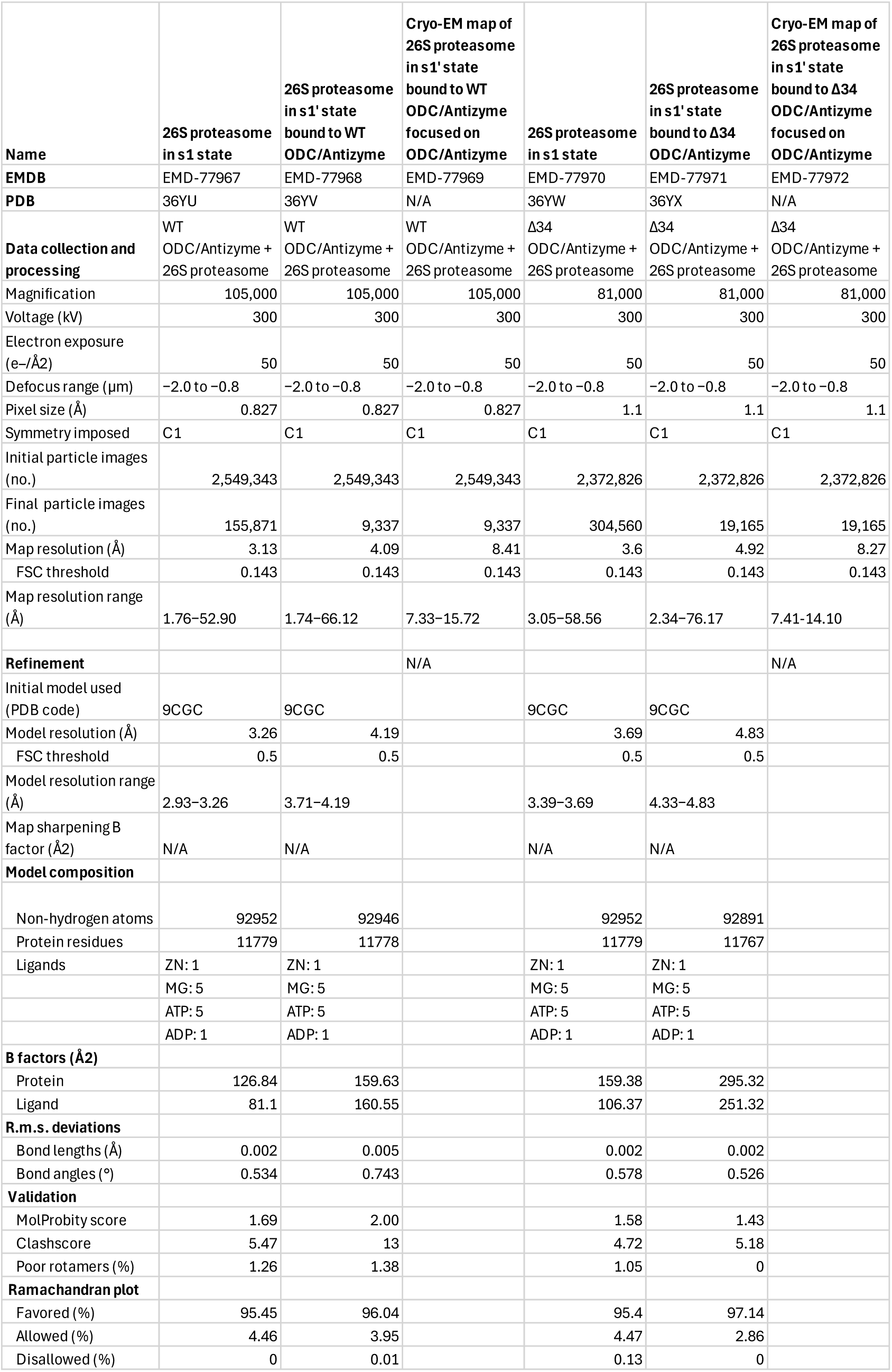
Cryo-EM data collection, refinement, and validation statistics.

To validate this interaction, we placed individual alanine substitutions in five conserved positions of yODC at the AlphaFold3-predicted interface with Rpt5 (Fig. 3f). The R338A, M346A, Y348A and W407A mutants of yODC showed diminished degradation in our SDS-PAGE-resolved end-point assay (Fig. 3g and Extended Data Fig. 3c), while the C411A mutation did not produce a statistically significant effect. These degradation reactions were performed at a sub-saturating substrate concentration (1 μM) and therefore reported on potential reductions in binding affinities. We corroborated this by a Michaelis-Menten analysis for the W407A mutant, which exhibited a 12-fold reduction in affinity (*K_M_* = 8.3 μM (95% CI [6.3, 11.5]) compared to *K_M_* = 654 nM (95% CI [480, 887]) for wild-type yODC; Extended Data Fig. 3d).

Docking the AlphaFold3 model for yODC/Antizyme:Rpt4/5cc into the composite map for the s1’ state 26S proteasome revealed that the first resolved residue in yODC’s N-terminal region, at position 35, is only 44 Å away from the entrance to 26S proteasome’s processing channel (Fig. 3c). A 34 amino-acid flexible initiation region can easily bridge this distance with sufficient extra length to enter the ATPase motor for engagement. In contrast, yODC’s C-terminus is located 84 Å away from the motor entrance, on the posterior side of yODC, which explains why transplanting the 34-residue flexible tail to the C-terminus did not allow degradation (Fig. 2e). Moving the first α-helix (35-58) as an additional spacer together with the flexible tail to the C-terminus remained unsuccessful for degradation as well, because it likely points in the opposite direction (Fig. 2e). Our structure shows how yODC’s N-terminus is ideally positioned for entering the proteasome’s processing channel, which also explains the strict requirement for an unstructured initiation region in this position.

### yODC and Antizyme form a two-part interface for proteasome binding

To assess how Antizyme allows yODC binding to the Rpt4/Rpt5 coiled coil, we compared the model of yODC/Antizyme:Rpt4/5cc with an AlphaFold3-prediction of the yODC homodimer (Fig. 3d-e). Interestingly, a large portion of yODC’s dimerization interface also serves as the binding surface for Antizyme and Rpt5 (Fig. 3e and Extended Data Fig. 3g-j). The yODC homodimerization interface is predicted to burry 3,002 Å^2^ of solvent-accessible surface area and can be mapped to 83 residues. On the other hand, yODC interactions with Rpt5 and Antizyme have a combined buried surface area of 2,379 Å^2^ and can be mapped to 59 residues, 52 of which (88%) are also predicted to be part of the homodimerization interface (Fig. 3e). This indicates a mechanism where Antizyme binding stabilizes yODC’s monomeric conformation and exposes its Rpt5-binding site, while also directly contributing itself to proteasome binding by interacting with Rpt4.

### yODC/Antizyme binding allosterically stabilizes a pre-initiation state

Compared to the s1 resting state of the proteasome, the yODC/Antizyme-bound s1’ conformation shows Rpn1 rotated counterclockwise by ∼ 90° around its Rpt1/Rpt2 coiled-coil anchor point on the ATPase motor (Fig. 4a). This feature is reminiscent of the E_B_ state observed for the substrate-engaged h26S proteasome^36^. To elucidate how yODC/Antizyme binding may drive this conformational change from s1 to s1’ and whether substrate engagement is required for it, we collected an additional cryo-EM dataset for the 26S proteasome bound to Δ34yODC/Antizyme, which lacks a flexible initiation region and therefore cannot be engaged and degraded. Remarkably, the proteasome was observed in the same s1’ conformation (Extended Data Fig. 3a-b, Extended Data Fig. 9, Table 1), suggesting that the transition from s1 to s1’ was allosterically triggered by Δ34yODC/Antizyme binding to the Rpt4/Rpt5 coiled coil and is independent of motor engagement (Extended Data Fig. 4.a-b). To understand how yODC/Antizyme binding and adopting the s1’ state affect the activity of the proteasome, we measured the ATPase-hydrolysis rates in the presence of saturating Δ34yODC/Antizyme, wild-type yODC/Antizyme, or the ubiquitinated Titin-I27^V15P^-23-K-35 substrate using a NADH-coupled assay (Fig. 4b and Extended Data Fig. 4c). For the ubiquitinated substrate, we observed a typical ∼2-fold stimulation of the proteasome’s ATPase activity, which is attributed to a shift in the conformational equilibrium from the non-hydrolyzing resting state (s1) to ATP-hydrolyzing processing states (non-s1) during active degradation^53^. Surprisingly, we detected a similar ∼2-fold stimulation also in the presence of Δ34yODC/Antizyme, even though this substrate cannot be engaged and degraded. This stimulation cannot be attributed to ATP hydrolysis events occurring in the s1’ state or during the transition from the s1 resting state, as both the s1 and s1’ conformations show the same nucleotide occupancy, with five ATPs and a single ADP bound to the Rpt6 subunit in the seam position of the staircase (Fig 4c and Extended Data Fig 4d). As the s1 resting state, the s1’ state thus appears inactive in ATP hydrolysis. However, we propose that the s1’ state upon yODC/Antizyme binding is more dynamic and facilitates the transitions to the processing non-s1 conformations. This model is further supported by the observation that the proteasome has an impressive 7.5-fold stimulation in ATPase activity while degrading wild-type yODC/Antizyme (Fig 4b and Extended Data Fig 4c). The stimulation likely originates from a combination of: (1) populating the s1’ state upon initial binding (as in the presence of Δ34yODC/Antizyme), (2) non-s1 conformations becoming stabilized by substrate engagement and degradation (as in the presence of the ubiquitinated substrate), and (3) a larger fraction of the proteasome pool being in the process of degradation due to efficient initiation of yODC/Antizyme degradation.

**Figure 4.**
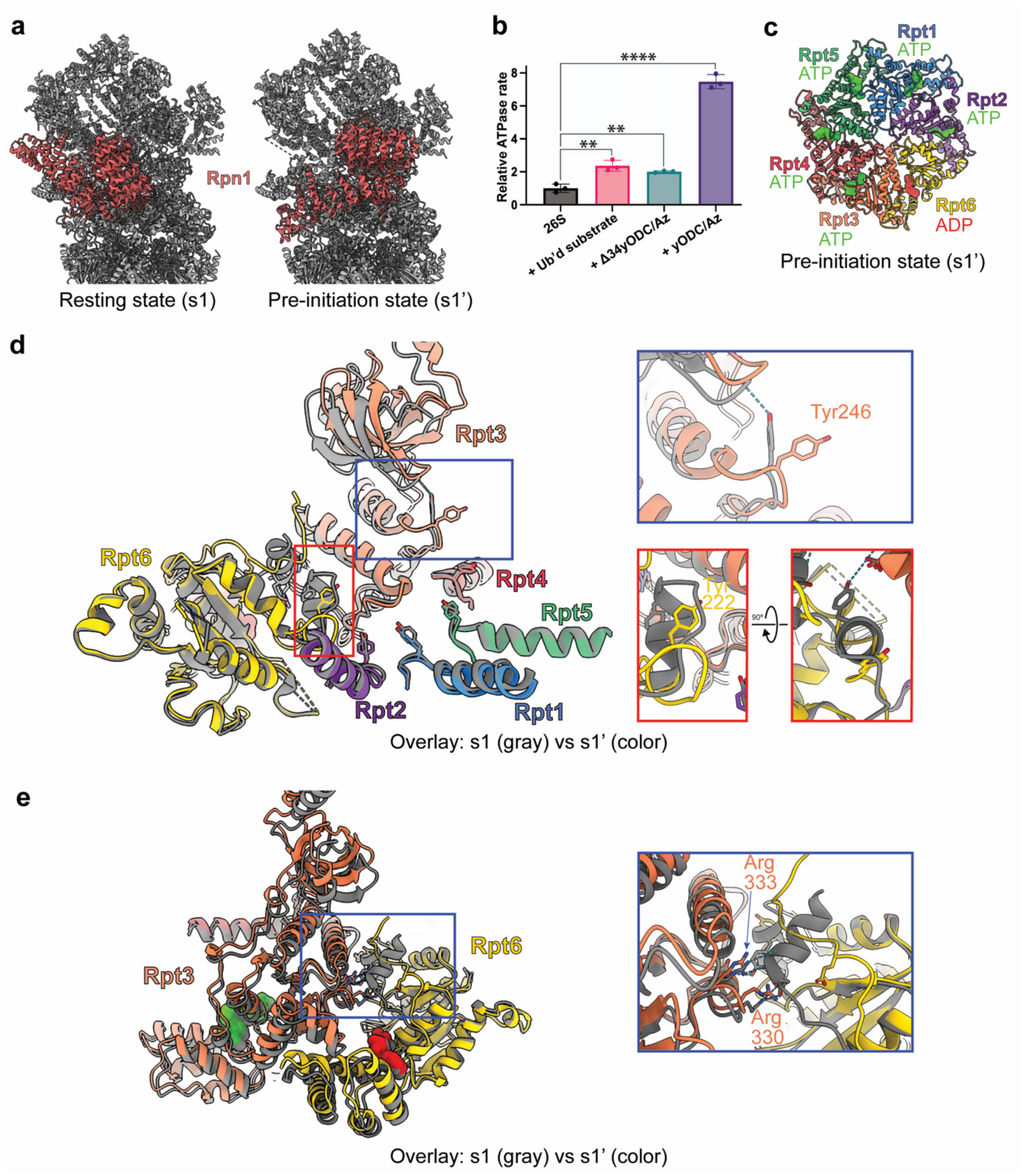
The proteasome transitions to a pre-initiation state prior to substrate engagement. **a**, Side-by-side view of the atomic models for the resting s1 state (left) and pre-initiation s1’ state (right), showing a rearrangement of ubiquitin-receptor subunit Rpn1 (red) upon yODC/Antizyme binding to the Rpt4/Rpt5 coiled coil. **b**, Normalized ATPase rates of the 26S proteasome in the presence of a ubiquitinated model substrate, yODC/Antizyme, or the N-terminally truncated Δ34yODC/Antizyme relative to the basal rate in the absence of any substrate. Shown are the mean values and standard deviations of the mean for n = 3 three technical replicates. Statistical significance was calculated using an ordinary one-way ANOVA with Dunnett’s multiple comparisons test against the proteasome-alone control (n = 3 technical replicates): **p = 0.0014 (+ubiquitinated substrate), **p = 0.0088 (+Δ34ODC/Az), and ****p < 0.0001 (+ODC/Az). **c**, Top-view of the ATPase hexamer in the s1’ conformation with the N-terminal domains of Rpt1-Rpt6 removed to show nucleotide occupancies, with ATP in green and ADP in red surface representation. **d**, Cutaway view of the central channel and the pore-1 loops with the conserved tyrosine for each Rpt subunit in stick representation. Shown is an overlay of the atomic models for the resting s1 (gray) and the pre-initiation s1’ (color) states in which Rpt3 is at the top and Rpt6 at the seam of the spiral staircase. The zoom-in views on the right highlight the rearrangements during the s1-to-s1’ transition, where Rpt3’s Tyr 246 loses its hydrogen bond with the N-domain and Rpt6’s Tyr222 get detached from Rpt3 as its pore-1 loop loses its helical conformation. **e**, View of the Rpt6/Rpt3 interface rotated 90°counterclockwise relative to the orientation in **d** to highlight Rpt3’s Arg fingers that in the s1 state stabilize the helical conformations of Rpt6’s pore-1 loop and the linker between the N-domain and ATPase domain. The zoom-in view on the right shows how these helices resolve and both Arg fingers are released during the s1-to-s1’ transition.

### Key contacts of the s1 resting state are dissolved in the s1’ pre-initiation state

To elucidate how the s1-to-s1’ transition may facilitate degradation initiation, we further examined the conformational rearrangements specific to the proteasome’s molecular motor. Typically, the asymmetric ATPase motor is arranged in a right-handed spiral-staircase conformation with one or two disengaged Rpt subunits at the seam position ^41,43^. While the spiral staircase in processing non-s1 states is dynamic with different Rpt subunits at the seam position, the staircase in the s1 resting state is static and apparently rigidified by interactions between the pore-1 loop tyrosine of Rpt1, Rpt2, and Rpt6 with Rpt3, and Rpt3’s tyrosine contacting the N-terminal domain ring^39^. Rpt6’s pore-1 loop thereby adopts a peculiar helical conformation that forms more extensive interactions with Rpt3 and abuts Rpt3’s arginine fingers, which normally coordinate the nucleotide bound to Rpt6 (Fig. 4d-e). Interestingly, the s1’ state in the Δ34yODC/Antizyme-bound proteasome shows Rpt3’s pore-1 loop tyrosine disengaged from the N-terminal domain and shifted towards the central channel (Fig. 4d and Extended Data Fig. 4e). Furthermore, Rpt6’s pore-1 loop no longer adopts an α-helical conformation, which moves the pore-1 loop tyrosine close to the central channel for substrate engagement and frees Rpt3’s arginine fingers to coordinate the Rpt6-bound nucleotide (Fig. 4f and Extended Data Fig 4e). Removing these s1 staples is thus expected to prime the ATPase motor for engaging an incoming substrate polypeptide and to facilitate the transition from the resting-conformation to the ATP-hydrolyzing processing states.

Comparing the s1’-state structures for the proteasome bound to the Δ34yODC/Antizyme versus wild-type yODC/Antizyme with the N-terminal tail inserted into the central channel showed virtually identical conformations of the ATPase motor (Extended Data Fig. 5). This suggests that s1’ conformation is indeed a pre-initiation state that is induced or transiently stabilized by yODC/Antizyme binding to the Rpt4/Rpt5 coiled coil, primes the motor for engagement of the flexible initiation region, and facilitates a timely transition to substrate-processing states.

### The s1’ pre-initiation state shows increased conformational dynamics

In order to further investigate the conformational effects of yODC/Antizyme binding on the dynamics of the s1’ state, we utilized a previously developed single-molecule FRET-based assay for the conformational transitions of the regulatory particle (Fig. 5a) ^24,39,40^. In this assay, the FRET-donor fluorophore LD555 is attached to an azido-phenylalanine (AzF) that replaces Phe2 in the lid subunit Rpn9, the FRET-acceptor fluorophore LD655 is attached to an AzF substituted for Ser26 in the N-terminal helix of Rpt5, and labeled proteasomes are immobilized on slides for TIRF (Total Internal Reflection Fluorescence) microscopy. Due to the strong structural similarities between the s1 state and the yODC/Antizyme-bound s1’ state, we do not expect a change in FRET efficiency for these two conformations. However, the switch from the s1 or s1’ to non-s1 states leads to a > 30 Å decrease in the distance between the fluorophore-attachment points and consequent increase in the apparent FRET efficiency^24,39,40^. Using a previously described Hidden Markov Model technique (ebFRET), we analyzed the dwell-time distributions for the low- and high-FRET states of proteasomes in the absence or presence of Δ34yODC/Antizyme (Fig. 5b-c; Extended Data Fig. 10). Δ34yODC/Antizyme causes a ∼ 2-fold acceleration of both the forward and backward conformational transitions, reflected in shorter dwell times (τ_s1-to-nons1_^ODC^ = 0.96 ± 0.11 s and τ_nons1-to-s1_^ODC^ = 0.47 ± 0.06 s versus τ_s1-to-nons1_ = 1.9 ± 0.2 s and τ_nons1-to-s1_ = 0.87 ± 0.09 s). Furthermore, we observed that the low- and high-FRET distributions are broadened and there is a small yet significant population of molecules with apparently continuous intermediate FRET efficiencies (Fig. 5c), which may be indicative of transient states. These findings suggests that the Δ34yODC/Antizyme-bound s1’ state is more dynamic and readily transitions toward non-s1 states, yet in the absence of an engageable initiation region also rapidly switches back. Interestingly, the faster s1-to-non-s1 transition, broader FRET distributions, and transient states with intermediate FRET efficiencies in the presence of Δ34yODC/Antizyme resemble a previously characterized proteasome variant with Tyr-to-Ala mutation in the pore-1 loop of Rpt6 (YA6)^39^. Since Rpt6’s pore-1 loop forms extensive interactions with the neighboring Rpt3 in the s1 state, removing the Tyr causes more frequent transitions to non-s1 states and correspondingly stimulates the basal ATPase rate by ∼ 2.5-fold, similar to what we observed for the s1’ state.

**Figure 5.**
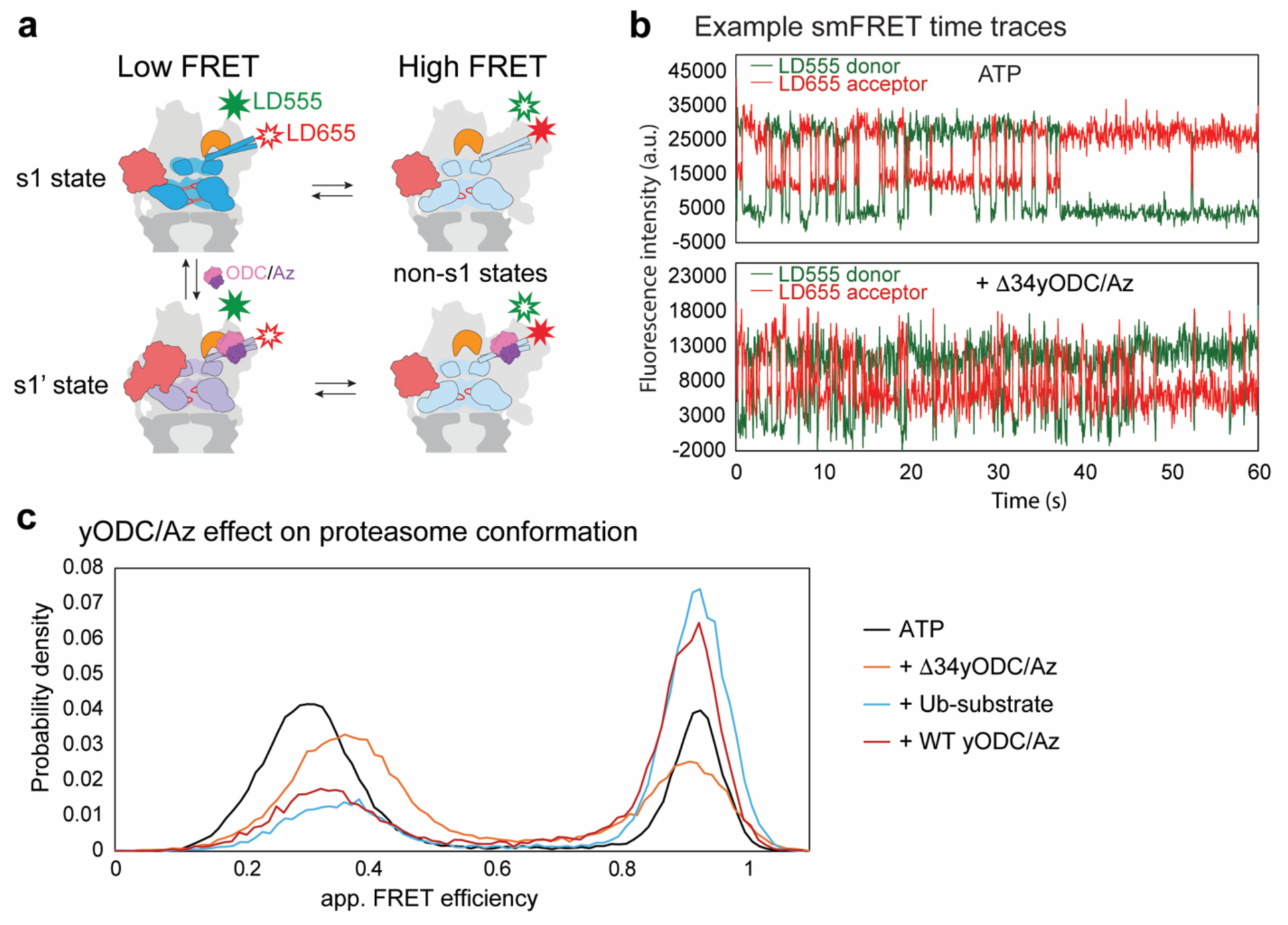
Binding the globular portion of yODC/Antizyme increases the 26S proteasome conformational dynamics. **a**, Schematic for the FRET-based assay to monitor the proteasome conformational transitions between the low-FRET resting s1 or pre-initiation s1’ states and the high-FRET non-s1 states. Proteasomes were labeled with a FRET-donor dye (LD555, green star) on the lid subunit Rpn9 and a FRET-acceptor (LD655, red) on the base ATPase subunit Rpt5. **b**, Example time traces for single-molecule TIRF measurements of the LD555 donor fluorophore attached to the lid subunit Rpn9 and the LD655 acceptor fluorophore attached to the base subunit Rpt5. Conformational switching leads to transitions between low FRET (s1 or s1’ state) and high FRET (non-s1 states). **c**, Single-molecule FRET distributions of the 26S proteasome in the absence or presence of Δ34yODC/Antizyme, a ubiquitinated titin^V15P^ model substrate, or WT yODC/Antizyme. Low FRET values represent the s1 and s1’ states, while high-FRET values are observed for non-s1 states ^39,40^. Number of single molecules used for analysis: 93 for ATP, 73 for Δ34yODC/Antizyme, 40 for Ub-substrate, and 34 for WT yODC/Antizyme.

Notably, our original FRET-based assay used an AzF-attached acceptor fluorophore at position 49 of Rpt5. However, with this proteasome construct we observed no yODC-Antizyme-induced changes in conformational transitions, likely because Gln49 is located at the center of the yODC binding site on the Rpt5 N-terminal helix (Fig. 3f), which independently supports our structural models. These single-molecule measurements therefore required a relocation of the fluorophore to position 26 in Rpt5.

## Discussion

More than 30 years ago, ODC was described as the first ubiquitin-independent substrate of the 26S proteasome, with Antizyme as a critical regulator^5-7^. Subsequent work revealed that the N-terminal region of yODC plays an essential role in facilitating degradation^12^, but the structural basis for Antizyme-mediated proteasome targeting and degradation of yODC remained unclear. In this study, we establish that the N-terminus of yODC is a sequence-independent initiation region that by itself is not sufficient for proteasome binding or degradation. Instead, yODC/Antizyme uses a bipartite degradation signal, similar to ubiquitinated substrates. The yODC/Antizyme complex is recruited to the 26S proteasome through interactions of its globular domains with the Rpt4/Rpt5 coiled coil, in a geometry that favors insertion of yODC’s 34-residue N-terminal initiation region into the molecular motor. Previous work has implicated the Rpt4/Rpt5 coiled coil as a ubiquitin-chain binding site^36,45–48^, and structural studies combined with single-molecule measurements indicated that the proximal, substrate-attached ubiquitin moiety interacts with the Rpn11 deubiquitinase prior to substrate insertion into the motor ^36,44^, which places this ubiquitin near the yODC/Antizyme binding site on the Rpt4/Rpt5 coiled coil. Systematic characterization of ubiquitinated model substrates also revealed that the ideal length for a flexible initiation region lies at ∼ 34 amino acids^22,23,25^. yODC/Antizyme thus uses similar principles and exploits the same proteasome features that underlie ubiquitin-dependent substrate turnover, which is supported by our observation that yODC/Antizyme and a ubiquitinated model substrate are degraded with equal priorities. Other ubiquitin-independent delivery systems have been shown to also utilize ubiquitin receptors and/or Rpn11 for the delivery of partially unstructured substrates to the proteasome, indicating a common requirement for a proteasome-binding element and an unstructured initiation region for engagement by the motor^3,4,8,29,30^.

Our findings provide a structural basis for the Antizyme-mediated activation of yODC for degradation by the 26S proteasome. Antizyme has been shown to inhibit ODC activity by disrupting the ODC homodimer and forming a 1:1 complex with ODC monomers^11^. Furthermore, published crystal structures of the hODC homodimer and the hODC/Antizyme complex explain the mutual exclusivity of these interactions^54–56^, which is in agreement with our AlphaFold3 predictions for the yeast orthologs. Here, we find for the yeast system that Antizyme binds directly to Rpt4 and acts as a “molecular glue” for stabilizing yODC’s interaction with Rpt5. As there is significant overlap between yODC’s dimerization interface and the surface that binds both Antizyme and Rpt5, only the Antizyme-bound yODC monomer is suspectable to degradation. Unlike other ubiquitin-independent delivery systems, this fascinating mechanism requires the substrate to participate in its own recruitment to the proteasome. Intriguingly, we discovered that the yODC/Antizyme complex cannot be generated by simple mixing of its components, but may form with the help of chaperones, upon unfolding by molecular motors like Cdc48/p97, or co-translationally, adding another layer of regulation to the yODC degradation process. The co-dependence of yODC and Antizyme for proteasome binding argues against Antizyme-mediated delivery as a more general mechanism for substrate turnover by the 26S proteasome and poses interesting questions about the evolutionary pressures that led to this dedicated degradation system for yODC.

In contrast to yODC, the unstructured initiation region of hODC is located at the C-terminus^50^, an attribute specific to vertebrate organisms in general. We show that this evolutionary specialization is substantial enough to prevent degradation of a “humanized” yODC with a C-terminally attached *H. sapiens’* initiation region. Upon recruitment of yODC/Antizyme to the Rpt4/Rpt5 coiled coil, yODC’s N-terminal initiation region is oriented right over the proteasome’s processing channel, and swapping its location to the C-terminus constitutes a significant geometric disruption. Recent work suggests that hODC/Antizyme predominantly binds to Rpn10 using a similar interface as yODC/Antizyme, but is rotated by 180° such that hODC’s C-terminus points toward the proteasome’s processing channel for insertion^8^. Additionally, we observed that h26S proteasome cannot degrade yODC/Antizyme, suggesting that its binding interface on the Rpt4/Rpt5 coiled coil is not conserved either (Extended Data Fig. 1f). These distinct binding modes explain the requirements for different locations of the flexible initiation regions, but the reasons for these divergent mechanisms of ubiquitin-independent, Antizyme-mediated degradation of ODC in lower and higher eukaryotes remains unknown.

The 26S proteasome relies on coordinated conformational rearrangements in its regulatory particle to allow selective substrate binding and efficient degradation^31–38^. The resting s1 state with its static spiral staircase of ATPase subunits is thereby ideally suited for substrate insertion into the central channel, before engagement by the motor triggers the conformational transition to processing non-s1 states for ATP-hydrolysis-driven hand-over-hand translocation, unfolding, and co-translocational deubiquitination^24^. However, the allosteric networks that drive and coordinate these events are unknown. Here, we observed the proteasome in a pre-initiation or primed (s1’) state that is induced or stabilized by yODC/Antizyme binding to the Rpt4/Rpt5 coiled coil prior to insertion of the initiation region into the central channel. In this s1’ conformation, the pore-1 loop interactions of Rpt6 and Rpt3 that stabilize the resting s1 state are dissolved, the conserved Tyr paddles of these subunits move closer to the central channel, and the ATPase motor becomes primed for substrate capture. The stimulated ATPase activity, increased conformational dynamics, and faster switching rates we observed for the proteasome in the presence of yODC/Antizyme suggests that the s1’ state more readily transitions to the ATP-hydrolyzing non-s1 states and is thus expected to facilitate substrate engagement and commitment to degradation (Fig. 6).

**Figure 6.**
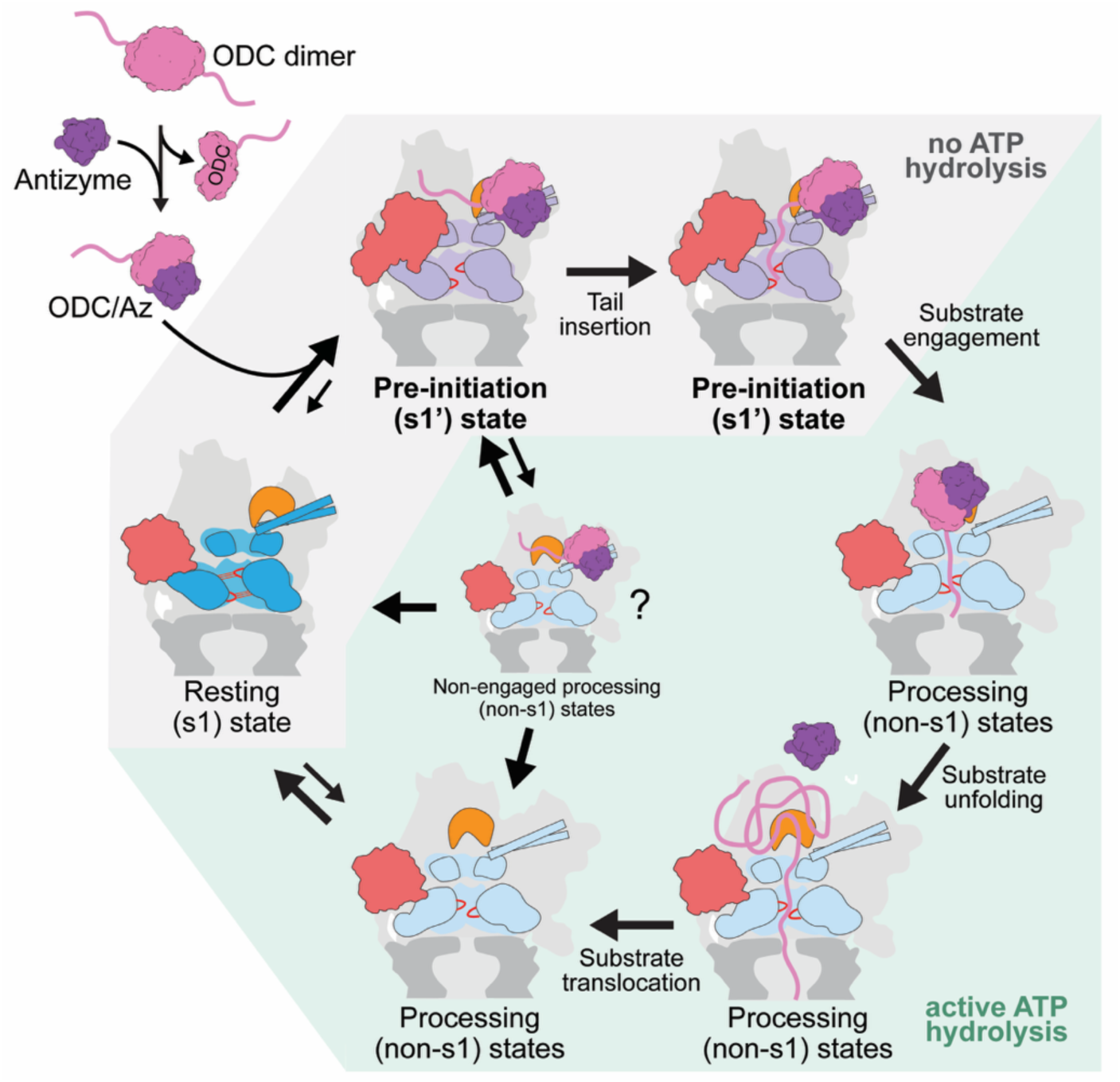
**Model for Antizyme-mediated delivery of yODC to the 26S proteasome. Top left**, Schematic for Antizyme-mediated separation of the ODC dimer and delivery of an Antizyme-bound monomer to the 26S proteasome. **Central cycle**, Binding of yODC/Az to the Rpt4/Rpt5 coiled coil of the resting (s1)-state proteasome stabilizes a pre-initiation (s1’) state, indicated by a large conformational change of Rpn1 (red). Insertion of yODC’s tail into the central channel leads to rapid engagement, switch to processing (non-s1) states, and ATP-dependent yODC unfolding, translocation, and degradation. The increased conformational dynamics of the yODC/Az-bound s1’ state may also lead to some sampling of the non-s1 states prior to yODC tail insertion, which is followed by either a rapid back-switching to the yODC/Az-bound s1’ state or yODC/Az dissociation and transition to the s1 state. Both the s1 and s1’ states have a static spiral staircase of ATPase subunits and are inactive in ATP-hydrolysis (gray underlay), but the s1 state is further rigidified through pore-loop interactions, indicated by red lines. ATP hydrolysis starts upon the substrate-induced or spontaneous switch to the processing non-s1 states with progressing spiral-staircase arrangements (green underlay).

We observed the same s1’ conformation after the N-terminal tail of yODC entered the central channel, suggesting that this state represents an on-pathway transition that likely aids degradation. That the pore-1 loop of Rpt6 dislodges from Rpt3 and moves closer to the central channel is thereby consistent with our recent biochemical and single-molecule studies that identified this pore loop as particularly important for initial substrate engagement and successful capture ^39^. Indeed, removing the Rpt6 pore-1 loop Tyr showed similar characteristics as the yODC/Antizyme-bound s1’ state, supporting our hypothesis that disengaging this loop from Rpt3 is a critical step in priming the ATPase motor. We propose that the Rpt4/Rpt5 coiled coil is an allosteric switch that detects the presence of a substrate and prepares the proteasome for substrate engagement. This also fundamentally agrees with our previous findings that ubiquitin-chain binding to Rpn11 in proximity to the Rpt5 coiled coil stabilizes the s1 state and helps substrate insertion into the motor^44^. Moreover, the s1’ conformation resembles the E_B_ state that was observed for the human 26S proteasome in the presence of a ubiquitinated substrate^36^. In this state, the substrate polypeptide was engaged by the ATPase motor, a ubiquitin moiety was bound to Rpn11’s catalytic groove, also contacting Rpt5, and the motor as well as Rpn1 showed arrangements similar to the s1’ conformation. The E_B_ state was suggested to represent a deubiquitination state of the regulatory particle, yet we propose that it may represent a pre-initiation state similar to s1’. Furthermore, we hypothesize that failure to insert a sufficiently long initiation region into the molecular motor while in the s1’ conformation results in a premature transition to non-s1 states and a rejection of the substrate (Fig. 6), which provides a potential model for the kinetic gateway that underlies the selection of ideal substrates^24^.

In summary, we biochemically and structurally characterized the degradation of a bona fide native substrate of the 26S proteasome. It establishes Antizyme’s role in making yODC accessible for degradation and acting as a molecular glue for yODC recruitment to the proteasome that allows subsequent engagement by the ATPase motor. Our results indicate that, similar to ubiquitinated substrates, ubiquitin-independent degradation systems rely on bipartite degradation signals, but use alternative strategies to satisfy these requirements. We found that yODC/Antizyme binding to a regulatory element in the Rpt4/Rpt5 coiled coil allosterically induces or stabilizes a primed conformation for efficient substrate engagement, and future studies will have to address whether this is a common strategy used by the 26S proteasome for selecting appropriate substrates in both ubiquitin-dependent and -independent manners.

## Methods

### Yeast strain and *Sc*26S proteasome purification

Endogenous Rpn11-3xFLAG-tagged 26S proteasome holoenzyme was purified from *S. cerevisiae* strain yAM3 (*MATa ade2-1 his3-11,15 leu2-3,112 trp1-1 ura3-1 can1-100 RPN11::RPN11-3XFLAG (HIS3) ubp6Δ::KANMX6*) as previously described^21^. Yeast cells were grown in YPD media at 30°C, harvested after four days and flash frozen on LN_2_. Frozen cells were then lysed cryogenically by grinding on a 6875 freezer mill dual chamber (SPEX Sample Prep). Lysates were resuspended in GF buffer (30 mM HEPES pH 7.6, 50 mM NaCl, 50 mM KCl, 5 mM MgCl₂, 5% glycerol) supplemented with 0.2% NP-40 and an ATP-regeneration system (5 mM ATP, 16 mM creatine phosphate, 0.03 mg/mL creatine kinase) and clarified by centrifugation at 26,000 rcf for 40 minutes at 4°C. Rpn11-3×FLAG tagged holoenzyme was affinity-purified from the clarified lysate using anti-FLAG M2 resin (ThermoFisher), washed with GF buffer containing 0.1% NP-40 and 0.5 mM ATP, and eluted with 0.3 mg/mL 3×FLAG peptide. The concentrated eluate was flowed over a Superose 6i 10/300 GL (Cytiva) using an ÄKTA pure chromatography system (Cytiva) in GF buffer supplemented with 0.5 mM ATP and 0.5 mM TCEP.

### Purification of h26S proteasomes

Human 26S proteasomes were purified from HEK293 cells stably expressing a C-terminal 6×His–TEV–biotin–6×His (HTBH) tag on Rpn11, originally generated by retroviral transduction and puromycin selection^57^ and adapted for suspension growth as previously described^4^. Cell cultures were maintained at 37 °C, 8% CO2, passaged twice weekly at ∼5 × 10^5^ cells/mL, with cryo-recovered cultures kept under puromycin selection. HTBH–Rpn11 HEK293 cells were grown from ∼5 × 10^5^ cells/mL for 72 h and harvested by centrifugation. Cell pellets were resuspended in lysis buffer (60 mM HEPES pH 7.4, 325 mM NaCl, 25 mM KCl, 5% glycerol, 10 mM MgCl2 and 0.5 mM TCEP) containing benzonase (Millipore), protease inhibitors (aprotinin, pepstatin, leupeptin and PMSF), 0.01% NP-40 and 2 mM ATP, lysed by Dounce homogenization and sonicated on ice. Lysate was clarified by centrifugation at 26,000 rcf for 30 minutes at 4°C and 26S proteasomes were bound to high-capacity streptavidin agarose (Pierce), washed with lysis buffer, and released by on-resin TEV cleavage (one hour at room temperature or overnight at 4 °C). The eluate was concentrated and resolved over a Superose 6i 10/300 GL (Cytiva) in GF buffer supplemented with 0.5 mM ATP and 0.5 mM TCEP.

### Recombinant expression and purification of ODC and Antizyme

Wild-type or mutant yODC and 6×His-tagged yAntizyme were recombinantly co-expressed in *E. coli* Rosetta(DE3) from T7-inducible plasmids, under kanamycin (50 μg/mL) and ampicillin (100 μg/mL) selection, respectively. To preserve yODC’s native N-terminus throughout the purification, yODC constructs were expressed as a Smt3 (SUMO) N-terminal fusion. Cells transformed with yODC and Antizyme expression plasmids were grown in auto-induction media for eight hours at 37°C and then at 18°C overnight before harvesting. Cell pellets were resuspended in NiA buffer (50 mM HEPES pH 7.6, 250 mM NaCl, 20 mM Imidazole) supplemented with 2 mg/mL lysozyme, benzonase, and protease inhibitors (aprotinin, pepstatin, leupeptin and PMSF), lysed by sonication on ice and clarified by centrifugation at 26,000 rcf for 30 minutes at 4°C. The ODC/Antizyme complex was batch bound for 45 minutes at 4°C to a HisPur Ni-NTA Resin (ThermoFisher) via the 6×His tag on Antizyme, washed on a gravity column with 10 CVs of NiA buffer, eluted with NiB buffer (50 mM HEPES pH 7.6, 250 mM NaCl, 250 mM Imidazole) and concentrated. Finally, Smt3-ODC was released by Ulp1 cleavage for 30 minutes at room temperature and ODC/Antizyme complexes were flowed over a Superdex 200 pg 16/600 (Cytiva) using an ÄKTA pure chromatography system (Cytiva) in GF buffer supplemented with 0.5 mM TCEP. This co-expressed material was used for all degradation, kinetic, and structural experiments unless individually purified ODC or Antizyme is explicitly indicated. To purify ODC homodimer, it was expressed as a 10×His-Smt3 N-terminal fusion and the protocol described in this section was followed. To purify Antizyme, the protocol in this section was followed but co-expression with ODC and the Ulp1 cleavage step were omitted.

### Site-directed mutagenesis and plasmid constructs

Point mutations at the Rpt4/Rpt5-binding interface (R338A, M346A, Y348A, W407A, C411A), the N-terminal deletion (Δ34yODC, lacking residues 1–34), N-terminal replacements (HA, cycB34), α-helix substitutions (35cycB58, 2xPro [E36P, L43P]), and the C-terminal tail-relocation constructs (swap-34, swap-cycB34, swap-Hx-34, swap-Hx-cycB34, swap-Hx-hsODC, swap-Hx-hsODC-CtoA) were generated by around-the-horn site-directed mutagenesis or Gibson assembly and verified by whole-plasmid sequencing. The cycB34 and 35cycB58 sequences were derived from *A. punctulata* cyclin B (cycB34: HTFNNENVSCRLGGAASIAVQAPAQHTFNNENVS, 35cycB58: RLGGAASIAVQAPAQHTFNNENVS). The *hODC* C-terminal sequences used were comprised of residues 425–461 with and without the C441A substitution.

### Purification of model substrates

The ubiquitinatable model substrate TitinI27-23-K-35, a single-domain substrate with one engineered lysine for ubiquitin-chain attachment and a 35-amino-acid unstructured initiation region, was generated as previously described^24^. The chimeric substrates cycB34-TitinI27 and ODC58-TitinI27 were generated by fusing the indicated sequence (the Cyclin B-derived 34-residue sequence, or ODC residues 1–58, respectively) to the N-terminus of an 18-K-21-TitinI27 substrate. Both chimeras were expressed as 10×His-Smt3 N-terminal fusions to protect their unstructured N-termini and purified following the same protocol used for 10×His-Smt3-ODC.

### Sortase-mediated fluorescent labeling

yODC, cycB34-TitinI27 and yODC58-TitinI27 were labeled with a Fluorescein (FAM)-conjugated peptide via a sortase A-mediated transpeptidation reaction^58^. The FAM-labeled peptide was generated by incubating 500 μM GGGSGCHHHHHH peptide (GenScript) with 600 μM Fluorescein-5-Maleimide (ThermoFisher) in GF buffer at room temperature for five minutes and quenching the reaction with 10 mM DTT. The constructs were designed with an LPETGG C-terminal sequence and were added at 100 μM to the quenched peptide reaction along with 20 μM sortase A and 10 mM CaCl_2_ for one hour at room temperature. Labeled substrates were isolated by Ni-NTA purification and resolved over a Superdex 200 Increase 10/300 GL or Superdex 75 Increase 10/300 GL column. To label ODC while in complex with Antizyme, Antizyme’s 6×His tag was cleaved using TEV prior to the labeling step. Labeling reactions were verified by SDS-PAGE (Extended Data Figs. 1b, 2c–d).

### *In vitro* ubiquitination of model substrates

TitinI27-23-K-35, cycB34-TitinI27 and ODC58-TitinI27 were ubiquitinated in vitro at the single engineered lysine to generate K63-linked polyubiquitin chains. Substrate (10 μM) was incubated with Uba1, Ubc1, and Rsp5 (2.5 μM each) in GF buffer with 10 mM ATP at room temperature, using the same enzyme system as previously described^24^; ubiquitin was added in 20 μM increments every 30 min, up to 120 μM total, over the 3 h reaction. Ubiquitination reactions were verified by SDS-PAGE (Extended Data Figs. 2e–f).

### Purification of sortase, Ulp1, and ubiquitination machinery

Sortase A, Ulp1, and the ubiquitination enzymes Uba1 (murine), Ubc1 (yeast), and Rsp5 (yeast) were each expressed as 6×His-tagged constructs in *E. coli* Rosetta(DE3) and purified individually using a similar Ni-NTA pulldown procedure described above for ODC/Antizyme: cells were grown in auto-induction media for eight hours at 37°C and then at 18°C overnight before harvesting, and cell pellets were resuspended in NiA buffer supplemented with lysozyme, benzonase, and protease inhibitors, lysed by sonication, and clarified by centrifugation. Clarified lysates were batch bound to HisPur Ni-NTA resin, washed with NiA buffer, and eluted with NiB buffer. The His tag on Sortase A was subsequently removed by TEV protease cleavage. All five proteins were further purified by size-exclusion chromatography on a Superdex 75 pg 16/600 column in GF buffer with 0.5 mM TCEP.

### Purification of ubiquitin

*S. cerevisiae* ubiquitin was expressed and purified following a previously established protocol^59^. Ubiquitin was expressed from an IPTG-inducible pET28a plasmid in Rosetta(DE3) *E. coli*. Cultures were grown in terrific broth at 37°C to an OD600 of 1.5–2.0, then induced with 0.5 mM IPTG and grown overnight at 18°C. Harvested cells were lysed by sonication in 50 mM Tris-HCl (pH 7.6) supplemented with lysozyme (2 mg/mL), benzonase, and protease inhibitors (aprotinin, pepstatin, leupeptin, PMSF), and the lysate was clarified by centrifugation. Perchloric acid was added to the clarified lysate (60% stock, 0.5% final) to precipitate contaminating proteins, and the resulting ubiquitin-containing supernatant was dialyzed overnight against 50 mM sodium acetate (pH 4.5). The dialyzed sample was loaded onto a 5 mL HiTrap SP FF cation-exchange column (Cytiva) and eluted with a linear NaCl gradient (0–0.5 M) in the same sodium acetate buffer. Peak fractions were pooled and flowed over a Superdex 75 pg 16/600 column in GF buffer.

### SDS-PAGE-based degradation assays

Degradation reactions contained 50 nM purified 26S proteasome and 4 μM substrate unless otherwise indicated in each figure legend. Assays were performed in GF buffer with 0.5 mM TCEP, 5 mM ATP, 0.5 mg/mL BSA, and an ATP-regeneration system (0.03 mg/mL creatine kinase and 16 mM creatine phosphate) at 30°C (unless otherwise noted). Where indicated, ATP and the ATP-regeneration system were replaced with 5 mM ATPγS, or reactions were supplemented with 100 μM bortezomib or 3 mM o-phenanthroline (oPA). Reactions were started by mixing solutions of 26S proteasome and substrate at 2x concentration and quenched at the indicated times with SDS sample buffer and resolved by SDS-PAGE. Gels were imaged on a ChemiDoc Imaging System (Bio-Rad) using Stain-Free fluorescence or were stained with Coomassie and then imaged. Gel quantification was performed by densitometry analysis using Image Lab 6.1 software (Bio-Rad). ODC band intensity was normalized to Antizyme band intensity in each well to correct for loading errors. Normalized ODC band intensity was graphed as a function of time, and initial rates were determined from the slope of a linear regression fit to the data points in the steady-state phase.

### Fluorescence polarization-based degradation kinetics

Degradation of FAM-labeled substrates was tracked in real time on a BMG Labtech CLARIOstar plate reader with MARS software by monitoring the loss of fluorescence polarization (excitation = 482 nm, emission = 530 nm) produced by proteolytic release of the dye-labeled fragment. Assays were performed in GF buffer with 0.5 mM TCEP, 5 mM ATP, 0.5 mg/mL BSA, and an ATP-regeneration system (0.03 mg/mL creatine kinase and 16 mM creatine phosphate) at 30°C; reactions were initiated by mixing a 2x solution of 26S proteasome with a 2x solution of substrate and competitive inhibitor (where applicable). Degradation reactions contained 50 nM purified 26S proteasome. For Michaelis-Menten analysis, initial rates measured across 100–4000 nM substrate were fit to the Michaelis-Menten equation in GraphPad Prism version 11 to determine K_m_ and k_cat_. Positive-control (PC) and negative-control (NC) samples were generated by pre-incubating 4000 nM labeled substrate with chymotrypsin protease or buffer, respectively, to define the FP assay’s dynamic range. For the competitive inhibition assays with ODC/Antizyme mutants (Fig. 1c), degradation of 4 μM FAM-labeled yODC/Antizyme was monitored in the presence of 12 μM dark yODC/Antizyme mutants, and the resulting decrease in initial rate was determined. For the head-to-head yODC/Antizyme versus ubiquitinated substrate competition assays, dark substrate was titrated from 0–10000 nM against a fixed concentration of 2 μM FAM-labeled substrate, and the resulting decrease in initial rate was determined as a function of inhibitor concentration.

### ATPase activity assays

26S proteasome ATPase activity was measured with an NADH-coupled assay in which ATP regeneration is coupled to NADH oxidation, monitored as a loss of absorbance at 340 nm on a BMG Labtech CLARIOstar plate reader with MARS software. Reactions contained 50 nM 26S proteasome and an ATPase mix (5 mM ATP, 3 U/mL pyruvate kinase (Sigma), 3 U/mL lactate dehydrogenase (Sigma), 1 mM NADH and 7.5 mM phosphoenolpyruvate) with saturating substrate (4 μM ubiquitinated TitinI27-23-K-35, or 10 μM WT or Δ34yODC/Antizyme). Rates were normalized to the basal, substrate-free ATPase rate of 26S proteasome.

### Liquid chromatography-mass spectrometry of degradation products

Following an end-point degradation reaction of co-expressed yODC/Antizyme by 26S proteasome, peptide products were isolated from full-length protein and 26S proteasome components using a centrifugal concentrator with a 3 kDa molecular-weight cutoff. Peptides in the flow-through were analyzed by LC-MS and mapped onto the yODC primary sequence. Peptide-length statistics (mean 9.4 residues, 95% CI [9.2, 9.7]; median 9; range 6–18) were calculated from 281 peptide species identified.

### Cryo-EM sample preparation and data collection

Cryo-EM samples were prepared by manually mixing 3 μM purified 26S proteasome with 12 μM WT yODC/Antizyme or Δ34yODC/Antizyme in GF buffer with 0.02% NP-40, 0.5 mM TCEP and 5mM ATP and incubating them for 45 seconds on ice, a strategy designed to suppress ODC degradation and stabilize proteasome-bound intermediates. Three μL of samples were applied to a glow-discharged (25mA for 25s) UltrAufoil R 2/2 200-mesh gold grid (Quantifoil) and plunge-frozen using a Vitrobot Mark IV (ThermoFisher) set to 0 pN blot force, 2.5 seconds blot time, 100% chamber humidity and 12°C.

Cryo-EM data for the WT yODC/Antizyme complex were collected at the Pacific Northwest Cryo-EM Center (PNCC) using SerialEM on a Titan Krios G1 (ThermoFisher) operated at 300 kV and equipped with a Gatan BioContinuum HD imaging filter and K3 direct electron detector at a nominal magnification of 105,000x (physical pixel size 0.827 Å, collected in super-resolution mode). Movies comprised 50 frames (3.3 seconds total exposure) for a total dose of 50 e⁻/Å² across a defocus range of −2.0 to −0.8 μm. A total of 16,433 movies were collected.

Cryo-EM data for the Δ34yODC/Antizyme complex were collected at the Stanford-SLAC cryoEM Center using EPU on a Titan Krios G3i (ThermoFisher) operated at 300 kV and equipped with a Gatan K3 detector behind a BioQuantum energy filter at a nominal magnification of 81,000x (physical pixel size 1.1 Å, collected in super-resolution mode). Movies comprised 50 frames (3.27 seconds total exposure) for a total dose of 50 e⁻/Å² across a defocus range of −2.0 to −0.8 μm. A total of 13,014 movies were collected.

### Cryo-EM data processing and model building

Cryo-EM data for both datasets were processed in cryoSPARC (versions 4.6–5.0) ^60^. Movies were motion-corrected and dose-weighted using patch motion correction (output Fourier-crop factor 0.5; maximum alignment resolution 5 Å), and defocus parameters were estimated using patch CTF estimation (fit resolution range 25–3 Å). Particles were picked with the blob picker function using circular blobs of 190 and 450 Å, and an elliptical blob of 190 × 450 Å. Particles were extracted using a box size of ∼900 Å. In both datasets, particles were Fourier cropped to speed up processing and were curated using 2D classification, multi-class ab-initio reconstruction and heterogeneous refinement to remove ice contamination, aggregates, and other low-quality picks, yielding final curated sets of 217,537 (WT) and 222,813 (Δ34) particles for downstream 3D processing.

For each dataset, the curated particle stack was refined to a consensus structure of the 26S proteasome using a homogeneous refinement job with C2 symmetry, followed by C2-symmetry expansion to treat the two regulatory particles of each doubly capped proteasome as independent particles; this effectively doubled the number of particles. The symmetry expanded particle stack was recentered around the regulatory particle using the volume alignment tool and refined to a consensus structure using a local refinement job masking the singly capped proteasome. The particles were subjected to 3D variability analysis^61^ followed by 3D Variability Display in cluster mode to eliminate proteasomes without regulatory particles and resolve conformational heterogeneity between the resting (s1-like) and processing (non-s1) states.

For the WT dataset, the s1-like particles (165,217 particles) were reextracted at the physical pixel size and again subjected to 3D variability analysis^61^ followed by 3D Variability Display in cluster mode to segregate particles in the s1’ conformation (9,338 particles) from the rest of the particles in the s1 state (155,879 particles). Next, we generated the final EM structures by locally refining the s1 and s1’ particle stacks independently using a mask of the regulatory particle. The local map of yODC/Antizyme bound to the Rpt4/Rpt5 coiled coil was generated by masking this region and running a local refinement on the s1’ particle stack.

For the Δ34yODC dataset, the s1-like particles (344,921 particles) were reextracted at the physical pixel size and an initial 3D variability analysis and cluster-mode display separated 304,560 particles assigned to the s1 conformation from a 40,361-particle cluster enriched in the s1’ conformation; this cluster was subjected to a second round of 3D variability analysis and cluster-mode display to isolate a purified s1’ population of 19,165 particles for final refinement. Here, we also ran a local refinement on the purified s1’ particle stack to generate a local map of yODC/Antizyme bound to the Rpt4/Rpt5 coiled coil by masking this region.

Final gold-standard Fourier shell correlation (GSFSC) resolutions, estimated at the 0.143 threshold using the auto-tightened solvent masks generated by cryoSPARC during refinement, were 3.13 Å (WT s1) and 4.09 Å (WT s1’), and 3.60 Å (Δ34 s1) and 4.92 Å (Δ34 s1’). The local maps of yODC/Antizyme bound to the Rpt4/Rpt5 coiled coil were resolved to a GSFSC resolution of 8.41 Å (WT) and 8.27 Å (Δ34).

Atomic models were built for the s1 and s1’ states of both the WT and Δ34yODC/Antizyme datasets. For each reconstruction, a previously determined s1 26S proteasome structure (PDB ID: 9CGC) was rigid-body docked into the cryo-EM map as a starting model, and the fit was improved through iterative cycles of flexible fitting in ISOLDE^62^ and real-space refinement in PHENIX^63^. The AlphaFold3 model of yODC/Antizyme bound to the Rpt4/Rpt5 coiled coil was rigid body docked into the local maps (8.41 Å, WT; 8.27 Å, Δ34) but was not further refined, consistent with their resolution.

### Bioinformatic sequence analysis

Multiple sequence alignments of yODC and hODC were generated from the top 20 hits of a standard protein BLAST search against the NCBI refseq_protein (yODC) or swissprot (yODC and hODC) databases, then aligned and colored using the Clustal color scheme in Jalview.

### Cloning of Rpt subunits for single-molecule measurements

New position for single molecule probe labeling for base compatible with yODC/Antizyme binding was designed using a previously described plasmid^39^. Briefly, all six WT Rpts were combined into a pCOLADuet-1 vector, with each ORF having its own T7 promotor. An amber codon was introduced replacing Ser52 of Rpt5.

### Purification and labeling of recombinant base and lid subcomplexes

Recombinant subcomplexes were expressed, purified, and labeled using a previously described protocol^39^. Briefly, for base expression, *E. coli* BL21(DE3)* cells were transformed with pAM81, pAM83, pAM87, and the new Rpt plasmid described above. The transformed cells were grown at 37 °C in 6 L of dYT to OD_600_=0.5, then 1 mM p-azidophenylalanine (AzF, Amatek Chemical) and 1 mM IPTG were added. The cells were induced for 5 hours at 30 °C and then further cultured at 16 °C overnight. Cells were harvested by centrifugation and then resuspended in NiA buffer (60 mM HEPES pH 7.60, 100 mM NaCl, 100 mM KCl, 10 mM MgCl_2_, 10% glycerol, and 25 mM imidazole) supplemented with protease inhibitors (aprotinin, leupeptin, pepstatin, and 4-(2-aminoethyl)benzenesulfonyl fluoride hydrochloride (AEBSF)), and benzonase. Cells were lysed by sonication and the lysate was clarified by centrifugation at 15,000 rpm for 30 min at 4 °C. The supernatant was loaded to a 5 mL HisTrap FF crude (Cytiva) column, washed with 10 CV NiA buffer, and eluted with NiA buffer + 250 mM imidazole. The eluant was then loaded onto 5 mL M2 Anti-FLAG affinity resin (Sigma-Aldrich), washed with 10 CV NiA buffer, and eluted with 15 mL NiA buffer + 0.5 mg/ mL 3×FLAG peptide. The FLAG eluent was concentrated to ∼ 500 µL with a 100 kDa molecular weight cut off (MWCO) concentrator (Millipore). For fluorophore labeling, 1 mM 5,5-dithiobis2-nitrobenzoic acid (DTNB, Sigma Aldrich) was added to the concentrated base and incubated for 10 min on ice to prevent off target labeling. Then, 100 μM dybenzocyclooctane (DBCO)-LD655 (Lumidyne Technologies) was added and the reaction progressed overnight at 4 °C supplemented with ATP Regeneration Mix (creatine phosphate (VWR) and creatine kinase (Sigma Aldrich)). The next day, the reaction was quenched by first adding 300 μM free AzF for 5 min and then 5 mM DTT for 30 min at 4 °C. The labeled subcomplex was further purified by size-exclusion chromatography (SEC) using a Superose 6 increase 10/300 column (Cytiva) in GF buffer (30 mM HEPES pH 7.60, 50 mM NaCl, 50 mM KCl, 10 mM MgCl_2_, 5% glycerol) + 0.5 mM TCEP and 1 mM ATP. The concentration of base subcomplex was determined by Bradford with a BSA (SigmaAldrich) standard curve, and the labeling efficiency was determined by quantification of the dye absorbance using a Nanodrop.

For lid expression, *E. coli* BL21(DE3)* cells were transformed with pAM80, pAM86, pAM87, and pAM90. The transformed cells were grown at 37 °C in 6 L of dYT to OD of 0.5, then 1 mM AzF (Amatek Chemical) and 1mM IPTG were added. The cells were induced for 5 hours at 30 °C and then further cultured at 16 °C overnight. Cells were harvested by centrifugation and then resuspended in NiA buffer supplemented with protease inhibitors (aprotinin, leupeptin, pepstatin, and AEBSF), and benzonase. Cells were lysed by sonication and the lysate was clarified by centrifugation at 15,000 rpm for 30 min at 4 °C. The supernatant was loaded to a 5 mL HisTrap FF crude (Cytiva) column, washed with 10 CV NiA buffer, and eluted with NiA buffer + 250 mM imidazole. The eluant was then loaded onto 10 mL amylose resin (NEB), washed with 10 CV NiA buffer, and eluted with 20 mL NiA buffer + 10 mM maltose. The eluent was concentrated to ∼ 500 µL with a 100 kDa molecular weight cut off (MWCO) concentrator (Millipore). For fluorophore labeling, 1 mM DTNB was added to the concentrated lid and incubated for 10 min on ice to prevent off target labeling. Then, 100 μM DBCO-LD555 (Lumidyne Technologies) was added and the reaction progressed overnight at 4 °C supplemented with HRV3C protease to cleave the MBP. The next day, the reaction was quenched by first adding 300 μM free AzF for 5 min and then 5 mM DTT for 30 min at 4 °C. The labeled subcomplex was further purified by SEC using a Superose 6 increase 10/300 column in GF buffer + 0.5 mM TCEP and 1 mM ATP. The concentration of lid subcomplex as well as the dye labeling efficiency were determined by absorbance at 280 nm and 555 nm, respectively, using a Nanodrop.

### Single molecule fluorescence microscopy

The conformational dynamics of the proteasome was monitored using a previously described single molecule setup^39^. Flow chambers were built on the PEGylated and biotin modified glass coverslip, and neutravidin was immobilized on the coverslip through the biotin. For production of biotinylated CP, the yeast strain (yAM80) containing endogenous Pre1 C-terminally tagged with Avi-tag and Flag tag was grown in YPD at 30°C for 1 day until saturation. The cells were collected by centrifugation, resuspended in lysis buffer lysis buffer (60 mM HEPES pH 7.6, 500 mM NaCl, 1 mM EDTA, and 0.2% NP40), frozen as droplets in liquid nitrogen, and then lysed using a Cryomill 6875D (SPEX SamplePrep). The frozen lysate was thawed and clarified by centrifugation, and CP was affinity-purified using anti-Flag M2 affinity resin (Sigma-Aldrich). The FLAG eluent was biotinylated by incubating with 25 μM *E. coli* biotin ligase BirA,100 μM biotin, 10 mM ATP, and 10 mM MgCl_2_ in lysis buffer overnight at 4°C. The biotinylated CP was further purified using size exclusion chromatography in GF buffer (30 mM HEPES pH 7.6, 50 mM NaCl, 50 mM KCl, 10 mM MgCl_2_, 5% glycerol, and 0.5 mM TCEP) on a Superose 6 Increase 10/300 column (Cytiva). Proteasomes were reconstituted by combining 500 nM biotinylated core, 400 nM LD655-base, 600 nM LD555-lid, and 1 μM Rpn10 in assay buffer (GF buffer + 0.3 μM Rpn10, 6 mM Trolox, 0.5 mg/mL BSA, 2 mM ATP, and ATP regeneration system) for 10 min at room temperature. The reconstituted proteasome was diluted in assay buffer + PCA/PCD oxygen scavenging system to ∼ 100 pM based on the concentration of base and flowed onto the coverslip to be immobilized for 5 min. Different experimental conditions were created by washing the flow chamber with assay buffer + PCA/PCD + nothing (ATP), + 5 µM ubiquitinated titin^V15P^ (Ub-substrate), + 23 μM WT yODC/Antizyme, or + 24 μM Δ34yODC/Antizyme. The fluorescence microscopy was started right after the wash. Briefly, we used a Nikon Ti2 TIRF microscope setup with Nikon LUN-F laser box, 60× oil objective (1.49 NA, Nikon), perfect focus system (Nikon), and filter cube TRF59907 (Chroma). Movies were taken at 20 frames per s for 60 s on an iXon Ultra 897 (Andor) EMCCD camera with 532 nm laser TIR excitation. The fluorescent signal was split into green and red channels using a Hamamatsu W-view Gemini equipped with a T640lpxr-UF2 dichroic mirror (Chroma) and ET595/50-m and ET655lp bandpass filters (Chroma). The movies were analyzed using iSMS software suite. Briefly, colocalized green and red fluorescence spots showing anticorrelation in green and red channels were picked from the movie. The fluorescence time traces were manually inspected to discard photobleached or photoblinked region, and FRET histograms were constructed from those time traces. FRET transitions were analyzed using the hidden Markov model modality in iSMS to extract dwell time. We defined FRET states higher than 0.6 to be non-s1 and lower than 0.6 to be s1. The dwell time distributions showed deviation from single exponential distribution, so we determined the estimate of lifetime of these dwell events using the mean of all observed dwell time: 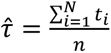, where *_t_*_$_ is the i-th observed dwell time and n is the number of observed dwell time. The standard error of the lifetime is then given by: 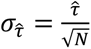, where N is the number of molecules used for dwell time analysis. The values of lifetime reported in main text were written as 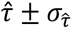.

## Resource availability

### Lead contact

Further information and requests for resources and reagents should be directed to and will be fulfilled by the lead contact, Andreas Martin.

### Materials availability

All constructs generated in this study are available from the lead contact upon request and completion of a Material Transfer Agreement.

### Data availability

· All data generated or analyzed during this study are included in this manuscript and the Supplementary materials. The cryo-EM density maps and corresponding atomic coordinates for the yeast 26S proteasome in the presence of yODC/Antizyme or its variant can be found on the Electron Microscopy Data Bank (EMDB) and Protein Data Bank (PDB) under the following accession codes: EMD-77967/PDB-36YU (s1-state 26S proteasome + WT yODC/Az), EMD-77968/PDB-36YV (s1’-state 26S proteasome + WT yODC/Az), EMD-77969 (s1’-state 26S proteasome + WT yODC/Az focused on yODC/Az), EMD-77970/PDB-36YM (s1-state 26S proteasome + Δ36yODC/Az), EMD-77971/PDB-36YX (s1’-state 26S proteasome + Δ36yODC/Az), EMD-77972 (s1’-state 26S proteasome + Δ36yODC/Az focused on Δ36yODC/Az).

- This paper does not report original code.
- Any additional information required to reanalyze the data reported in this paper is available from the lead contact upon request.

## Acknowledgments

We thank all members of the Martin lab for discussion and support. Cryo-EM data were collected at the Pacific Northwest Cryo-EM Center (PNCC) with assistance from Sean Mulligan and at the Stanford – SLAC Cryo-EM Center (S^2^C^2^) with assistance from Alexandre Cassago. We thank Robert Maxwell and the Vincent J. Coates Proteomics/Mass Spectrometry Laboratory Core Facility in the QB3 Institute for mass-spec data acquisition and help with analysis.

## Author contributions

D.R.R.-O., K.C.D., and A.M. conceived the study and designed experiments, D.R.R.-O. cloned constructs, expressed, and purified proteins, and performed biochemical with help by K.C.D.. D.R.R.-O. also performed cryo-EM sample preparation and data processing, and generated atomic models based on cryo-EM maps. H.H.H. performed single-molecule experiments and data analyses. D.R.R.-O. and A.M. wrote the manuscript.

---

**Extended Data Fig 1.**
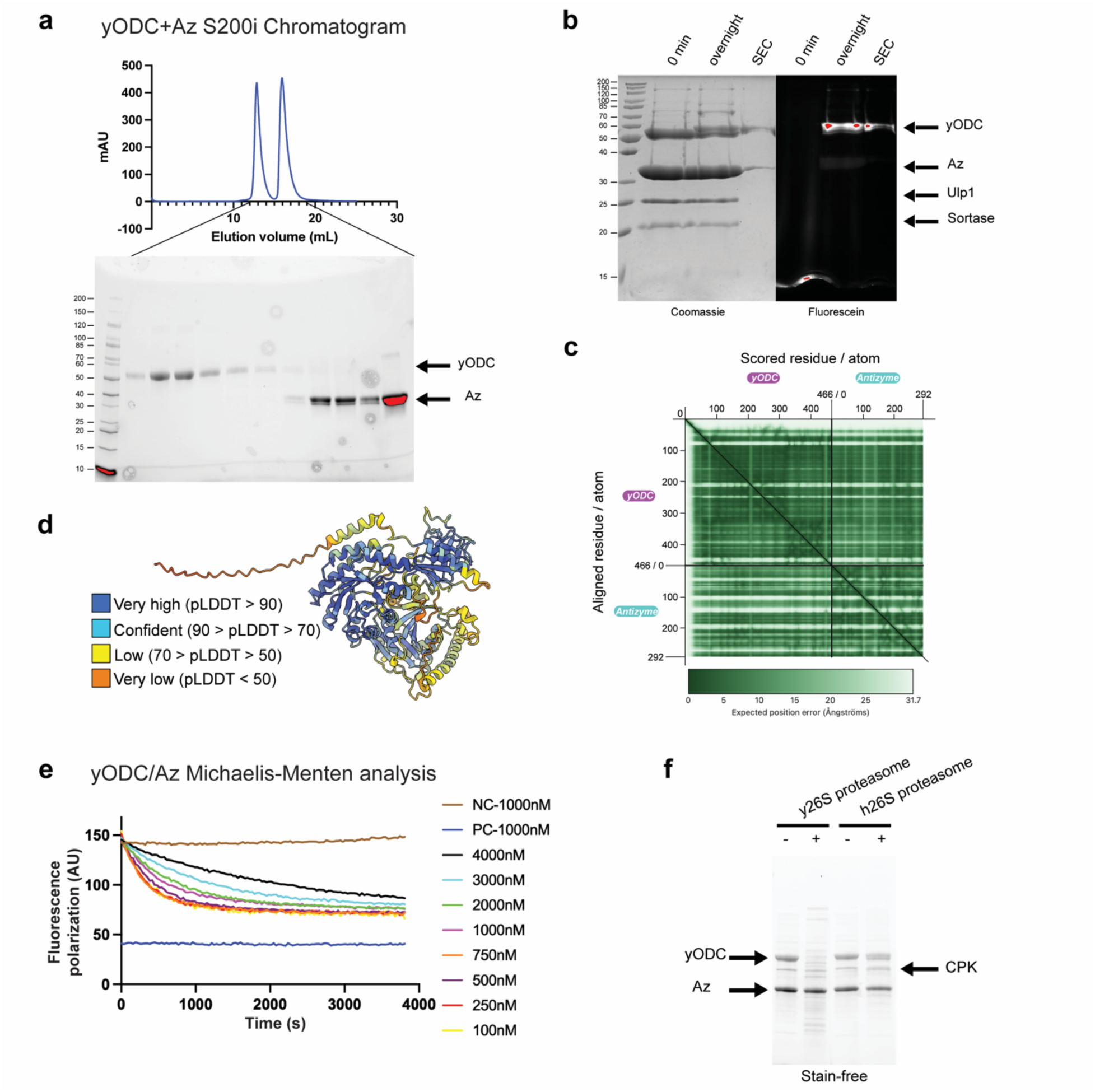
Supporting data for. Fig. 1**. a**, Top, Chromatogram for the Size Exclusion Chromatography (SEC) ofy ODC and Antizyme samples incubated for one hour. Bottom, SDS-PAGE analysis of SEC fractions. **b**, SDS-PAGE analysis of a sortase-catalyzed labeling reaction of yODC with FAM-modified peptide. **c**, Predicted Aligned Error (PAE) plot for an AlphaFold3-pedicted model of yODC/Antizyme. **d**, AlphaFold3-predicted model of yODC/Antizyme colored by predicted local distance difference test (pLDDT). **e**, Representative raw data from the Michaelis-Menten analysis of yODC/Antizyme degradation by the 26S proteasome. Initial degradation rates at various yODC/Antizyme concentrations (100-4000 nM) were linear fitting the decrease in FAM fluorescence polarization. The positive control (PC) and negative control (NC) samples were prepared by preincubating 4000 nM labeled yODC/Antizyme with chymotrypsin or buffer, respectively. **f**, SDS-PAGE analysis of yODC/Antizyme degradation by the yeast or human 26S proteasome. The 60-minute end points show that only y26S proteasome can degrade the yODC/Antizyme complex.

**Extended Data Fig 2.**
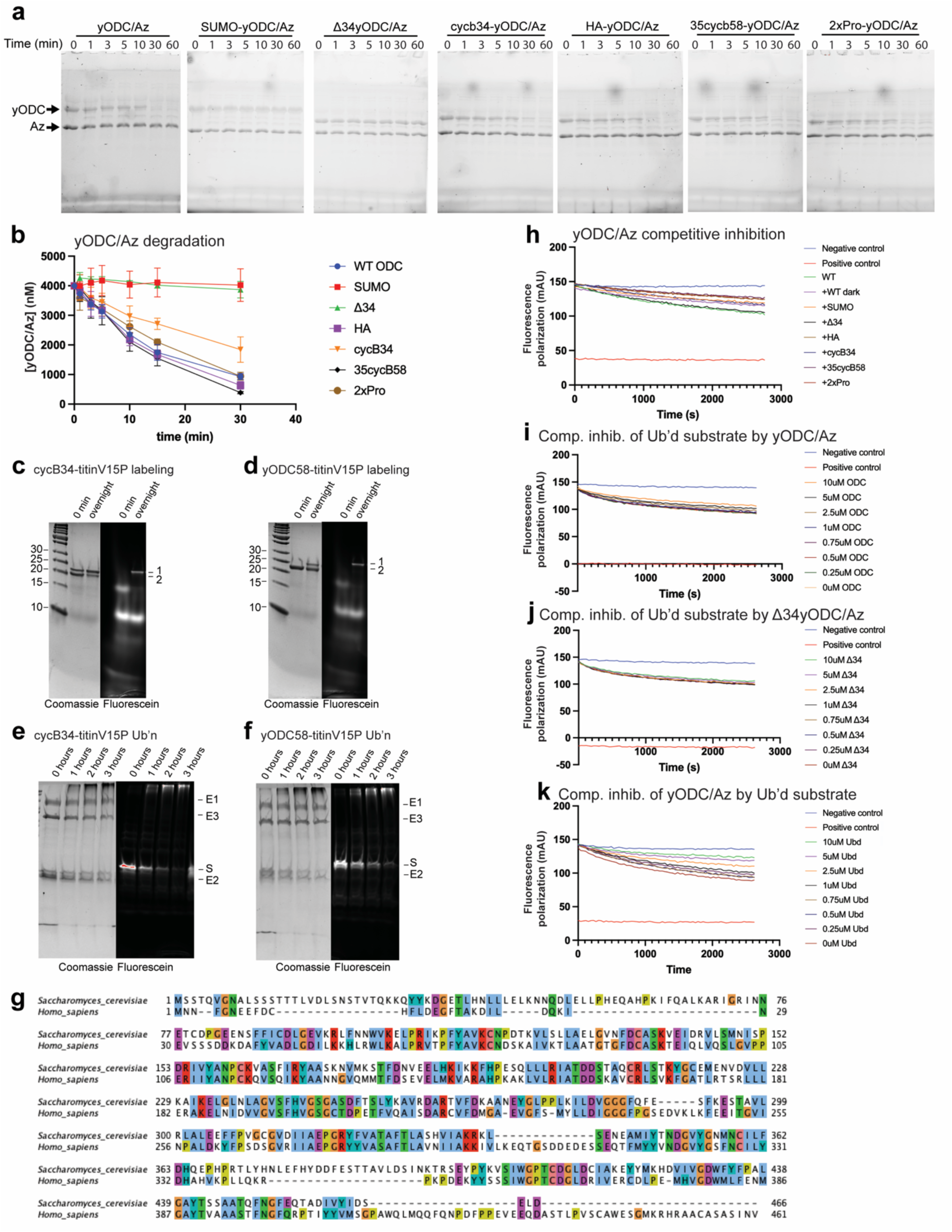
Supporting data for. Fig. 2**. a**, Representative time courses for the proteasomal degradation reactions of yODC/Antizyme mutants resolved by Coomassie-stained SDS PAGE. **b**, Plot of the average yODC/Antizyme amounts remaining as a function of time during proteasomal degradation, quantified from SDS-PAGE-resolved time-courses as shown in **a**. n = 3 technical replicates. **c-d**, SDS-PAGE analysis of sortase-labeling reactions for cycB34-titinV15P (**c**) and ODC58-titinV15P (**d**). **e-f**, SDS-PAGE analysis of the ubiquitination reaction for cycB34-titinV15P (**e**) and ODC58-titinV15P (**f**). **g**, Amino acid sequence alignment of *S. cerevisiae* and *H. sapiens* ODC colored using Clustal color scheme. **h**, Representative raw data for the competitive inhibition of labeled yODC/Antizyme degradation by unlabeled (dark) yODC/Antizyme variants (Fig. 2c). Initial rates for the degradation of 4 μM FAM-labeled yODC/Antizyme in the presence of 12 μM inhibitor were determined by linear fitting of the decay in FAM fluorescence polarization. **i**, Representative raw data from the competitive inhibition of Ub’d substrate degradation by yODC/Antizyme (Fig. 2d). **j**, Representative raw data from the competitive inhibition of Ub’d substrate degradation by Δ34yODC/Antizyme (Fig. 2d). **k**, Representative raw data from Competitive inhibition of ODC/Az by Ub’d substrate (Fig. 2e). **i-k**, Initial rates for the degradation of 4 μM FAM-labeled substrate in the presence of different concentration of inhibitor (0-10 μM) were determined by linear fitting of the decay in FAM fluorescence polarization. **h-k**, The positive control (PC) and negative control (NC) samples were prepared by preincubating 4000 nM labeled substrate with chymotrypsin or buffer, respectively.

**Extended Data Fig 3.**
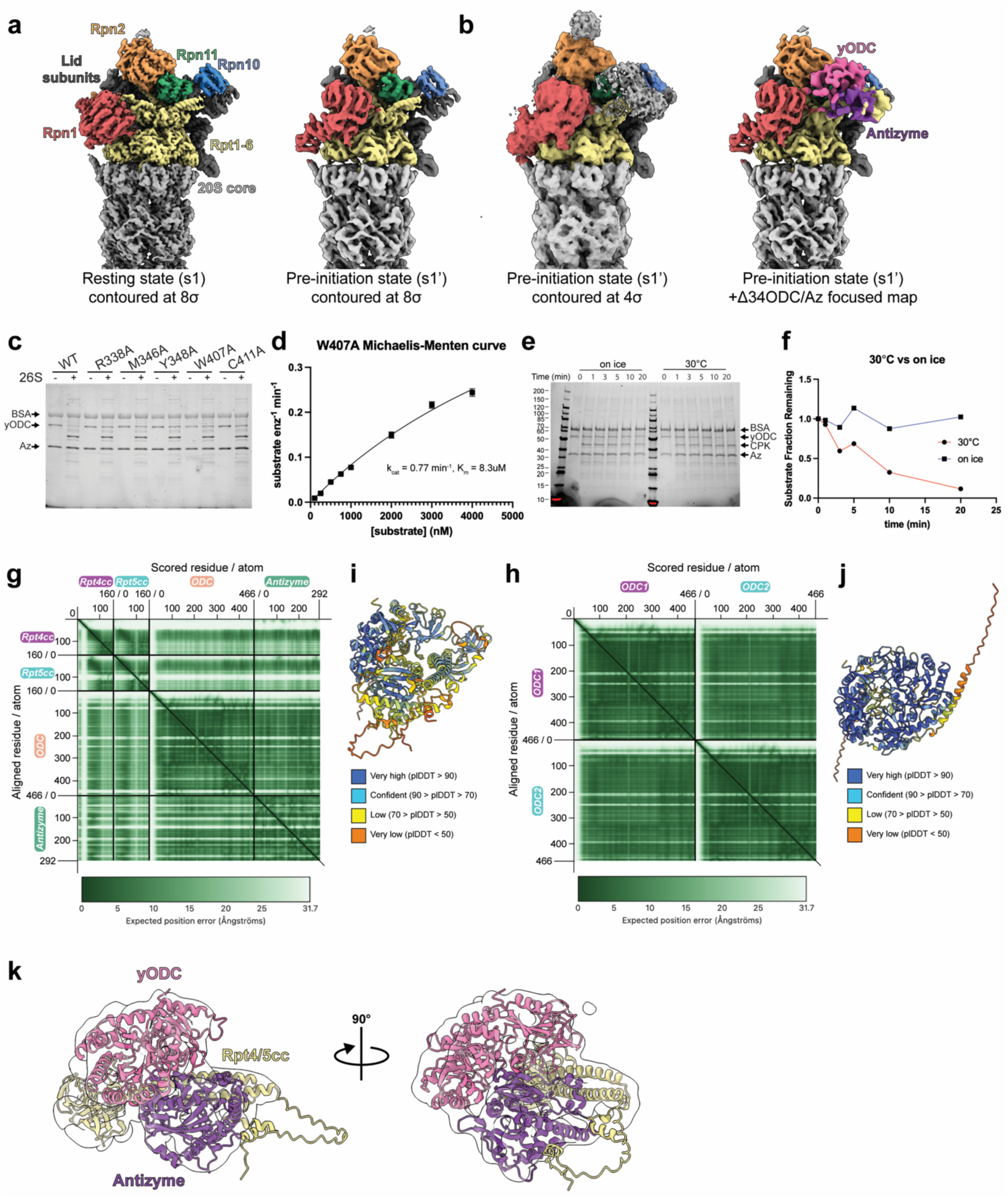
Supporting data for. Fig. 3**. a**, Cryo-EM density for resting state (s1) and pre-initiation state (s1’) proteasomes, colored by RP subunits. The proteasome was incubated with Δ34yODC/Antizyme on ice for 45 s before sample preparation. **b**, Left, Cryo-EM map of the s1’ state proteasome at low contour reveals unassigned density near the Rpt4/Rpt5 coiled coil. Right, focused cryo-EM map of additional density corresponding to the Δ34yODC/Antizyme complex overlayed onto the s1’ proteasome density. **c**, Degradation reactions of yODC/Antizyme binding mutants (1 μM) by 50 nM 26S proteasome, analyzed by Coomassie-stained SDS PAGE. The 7.5-minute end points show impaired degradation for most assayed mutants. **d**, Initial degradation rates for the W407A yODC/Antizyme binding mutant at various concentrations (100-4000 nM) were determined by linear fitting the decrease in FAM fluorescence polarization, and the resulting curve was fit to the Michaelis-Menten equation. **e**, SDS-PAGE-resolved time course for the degradation of 1 μM yODC/Antizyme by 50 nM 26S proteasome on ice vs. at 30°C. **f**, Plot of yODC/Antizyme amounts remaining as a function of time during proteasomal degradations shown in **e**. **g-h**, Predicted Aligned Error (PAE) plots for AlphaFold3-pedicted models of yODC/Antizyme bound to the Rpt4/Rpt5 coiled coil (**g**) and for the yODC homodimer (**h**). **i-j**, AlphaFold3-predicted model of yODC/Antizyme bound to the Rpt4/Rpt5 coiled coil (**i**) and the yODC homodimer (**j**) colored by predicted local distance difference test (pLDDT). **k**, AlphaFold3 model for yODC/Antizyme bound to the Rpt4/Rpt5 coiled coil is shown docked into the locally refined cryo-EM map of the yODC/Antizyme-bound proteasome. Amino acids 1-34 of ODC are not depicted.

**Extended Data Fig 4.**
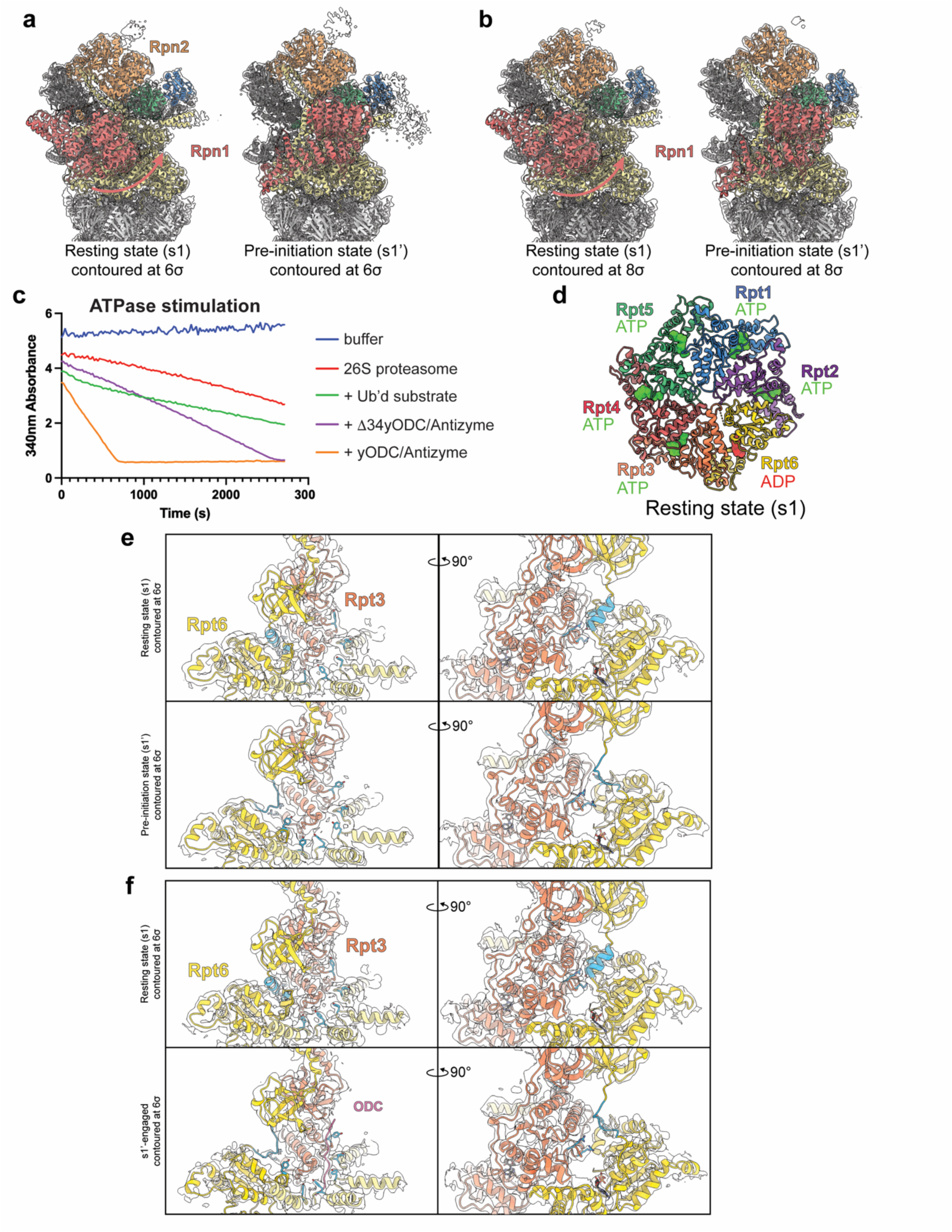
Supporting data for. Fig. 4**. a-b**, Side-by-side view of the atomic models docked into the cryo-EM density for the proteasome in the s1 state (left) and s1’ state (right), showing a rearrangement of ubiquitin-receptor Rpn1 (red, movement highlighted by arrow) upon yODC/Antizyme binding. Cryo-EM samples were prepared by incubating 26S proteasome with (**a**) WT yODC/Antizyme or (**b**) Δ34yODC/Antizyme on ice for 45 s before plunge-freezing. **c**, Representative raw data for the NADH-coupled ATP-hydrolysis assay (Fig. 4c) of proteasome in the presence of saturating concentrations of ubiquitinated substrate (4 μM), WT yODC/Antizyme (10 μM), or Δ34yODC/Antizyme (10 μM). **d**, Top-view of the ATPase hexamer in the resting s1 conformation with the N-terminal domains of Rpt1-Rpt6 removed to show nucleotide occupancies, with ATP in green and ADP in red surface representation. **e**-**f**, Cutaway view of the proteasome’s ATPase ring showing the top (Rpt3) and seam (Rpt6) subunits along with the other subunits’ pore-1 loops docked into Cryo-EM density. The pore-1 loop tyrosines, Rpt3’s arginine fingers, and an α-helix that is present in Rpt6’s linker between the N-terminal domain and ATPase domain when in the s1 state are colored sky blue.

**Extended Data Fig. 5.**
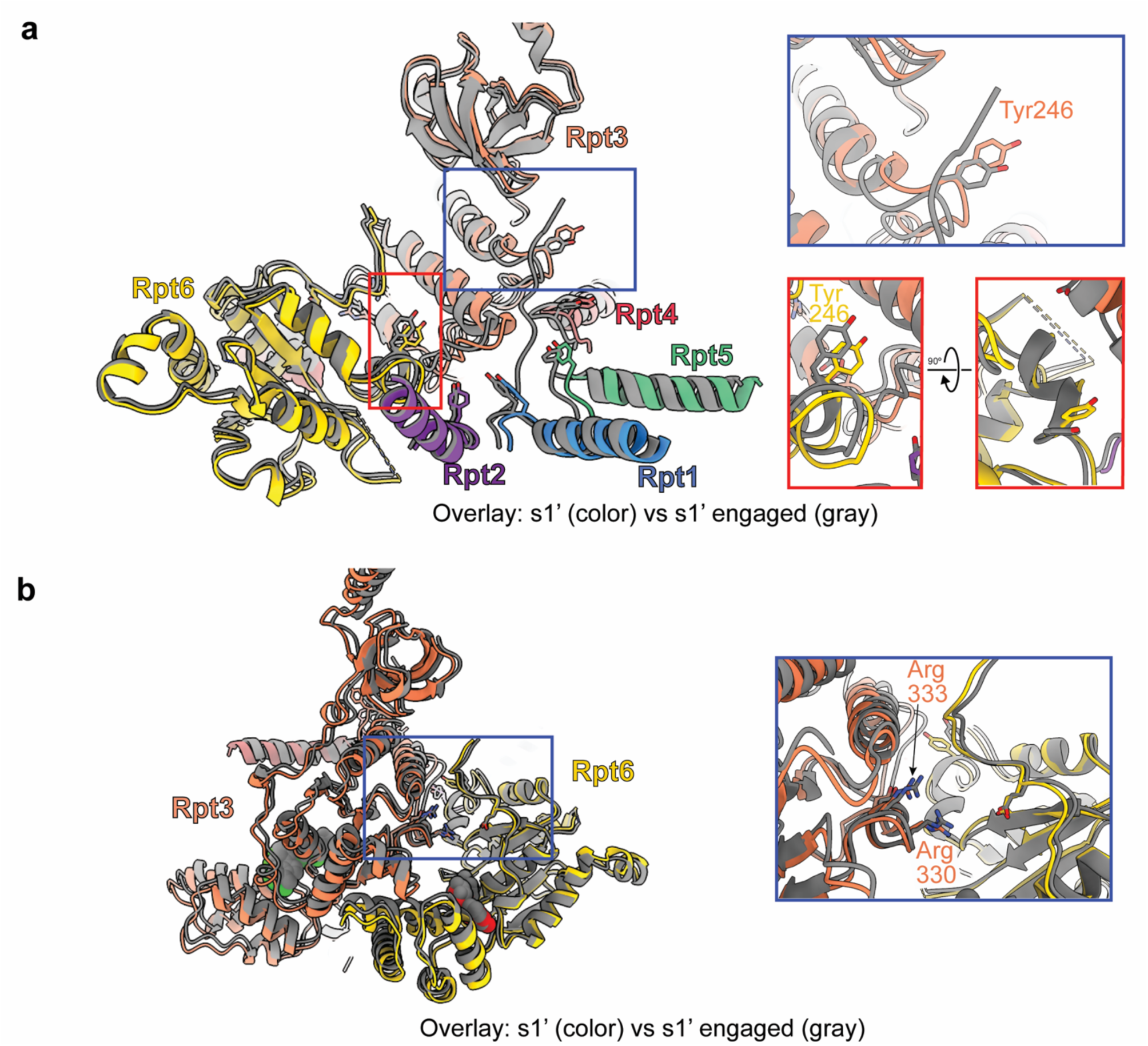
**The pre-initiation s1’ state shows similar conformations with and without inserted substrate tail. a-b**, Cutaway view of the central channel and pore-1 loops for the overlayed atomic models of the s1’ state without (color) and with inserted substrate tail (gray). Shown are the complete top subunit (Rpt3), the ATPase domains of the seam subunit (Rpt6), and the pore-1 loops of the 4 other Rpt subunits. The view in **b** is a counterclockwise 90°-rotation of **a**. The insets highlight the similar arrangement of the pore-1 loop tyrosines of Rpt3 and Rpt6 (**a**), and the released to arginine fingers of Rpt3.

**Extended Data Fig 6.**
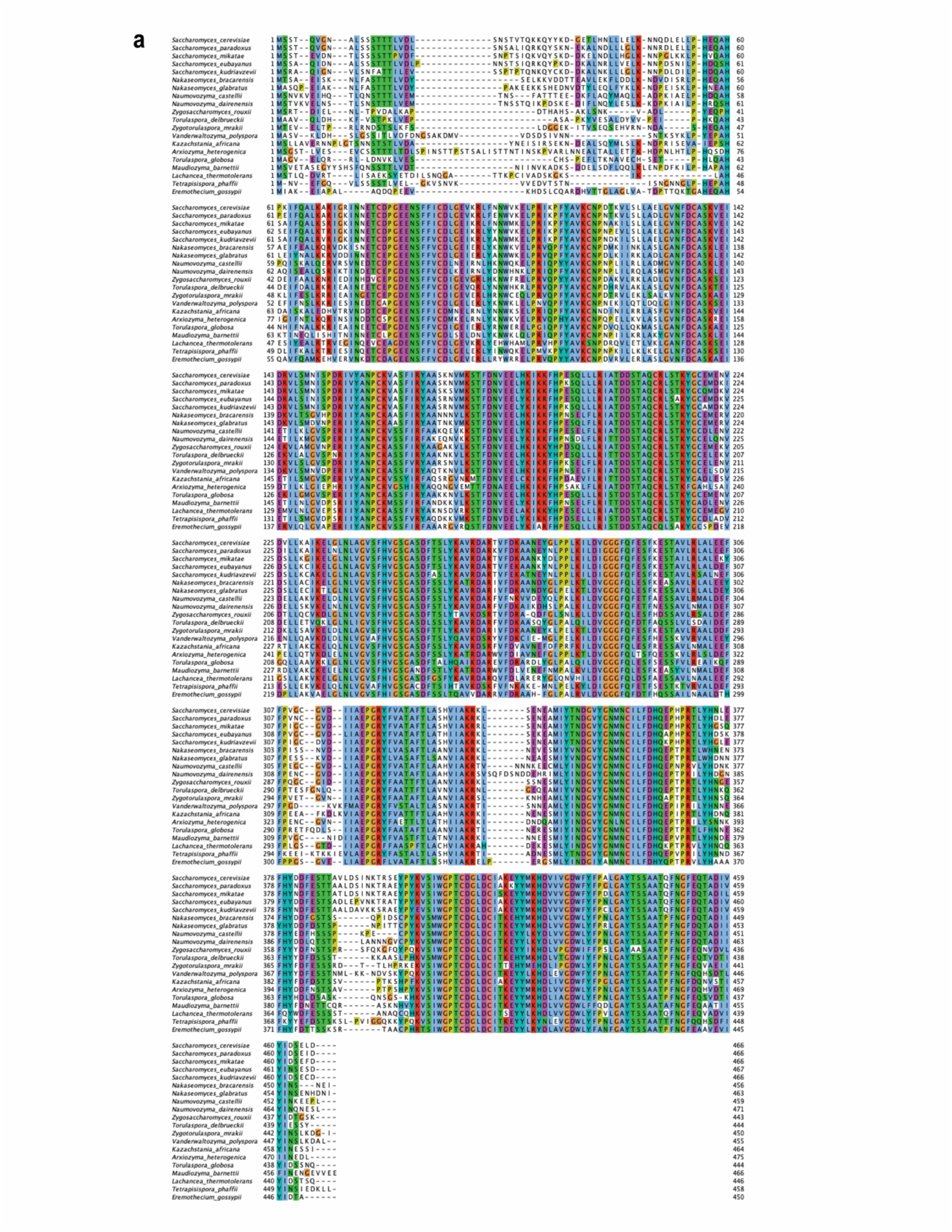
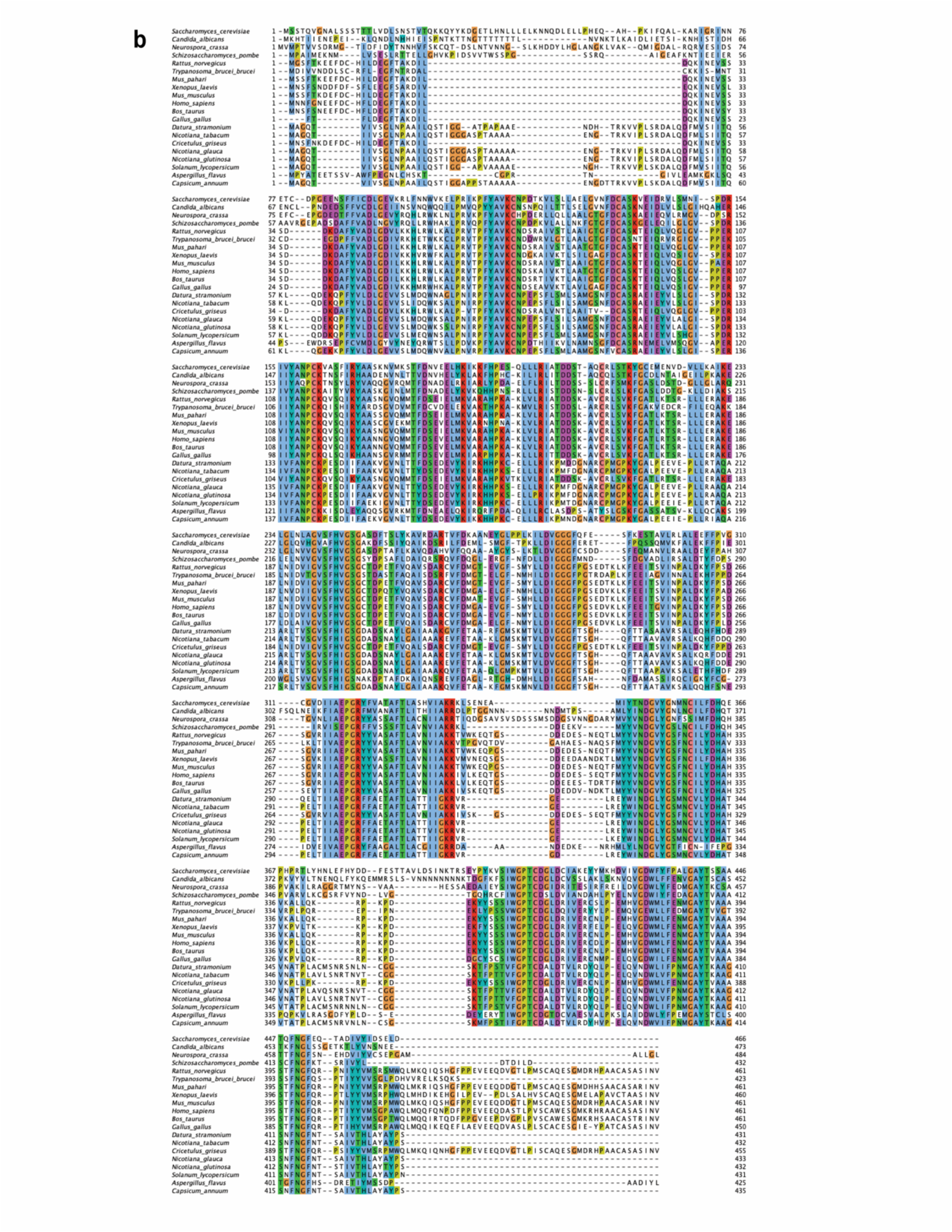
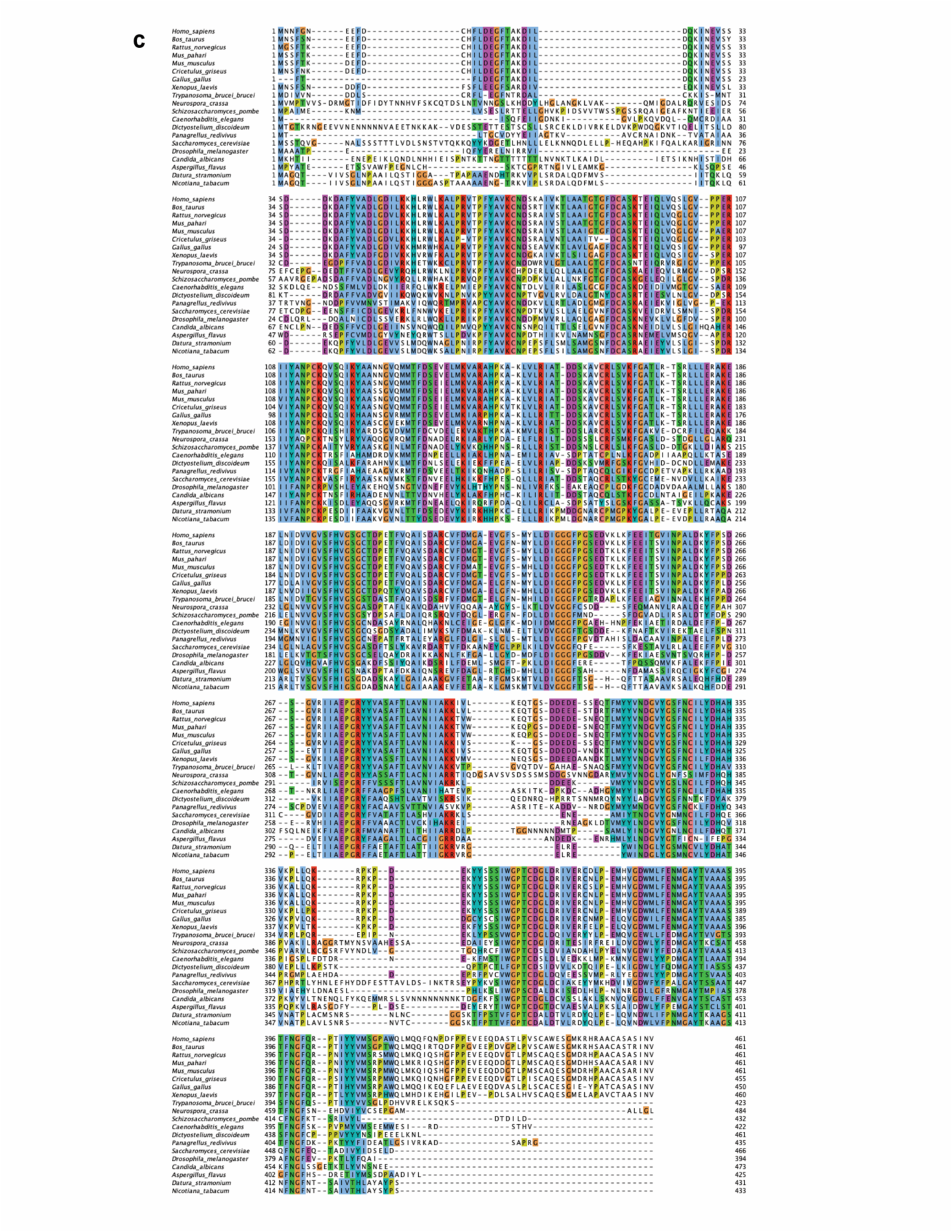
The N-terminal unstructured initiation region of yODC is not well conserved and vertebrate ODC has its initiation region at the C-terminus. a-c, Multiple sequence alignments (colored using the Clustal color scheme) of the top 20 hits from a query using standard protein BLAST against the stated NCBI database. **a**, query: yODC, database: refseq_protein. **b**, query: yODC, database: swissprot. **c**, query: hODC, database: swissprot.

**Extended Data Fig 7.**
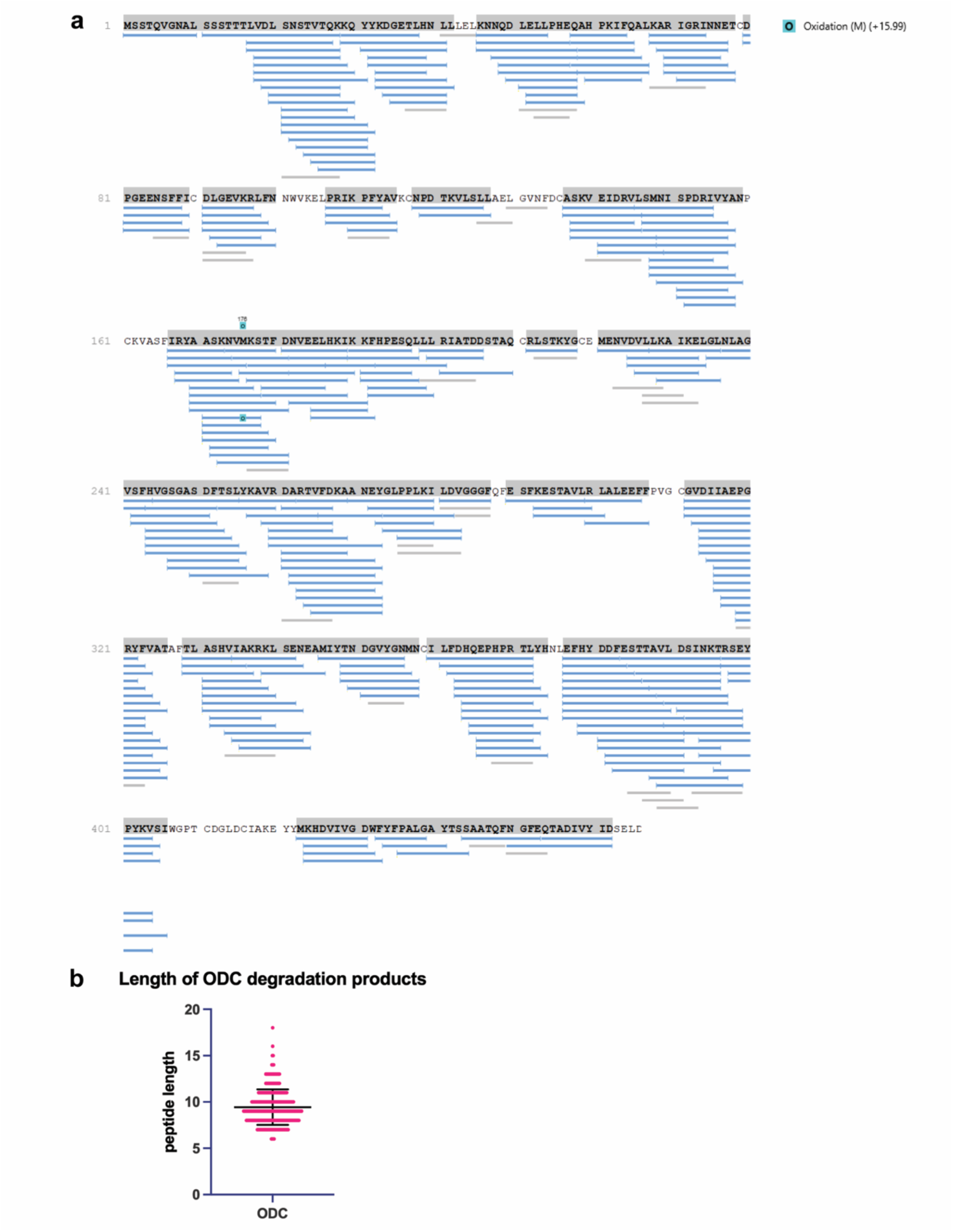
LC-MS analysis of yODC degradation products. **a**, Peptide products identified by LC-MS mapped onto the yODC amino acid sequence. **b**, Scatter plot of the amino acid lengths for yODC-peptide products. Mean: 9.4 (95% CI [9.2, 9.7]), Median: 9, Minimum: 6, Maximum: 18.

**Extended Data Fig 8.**
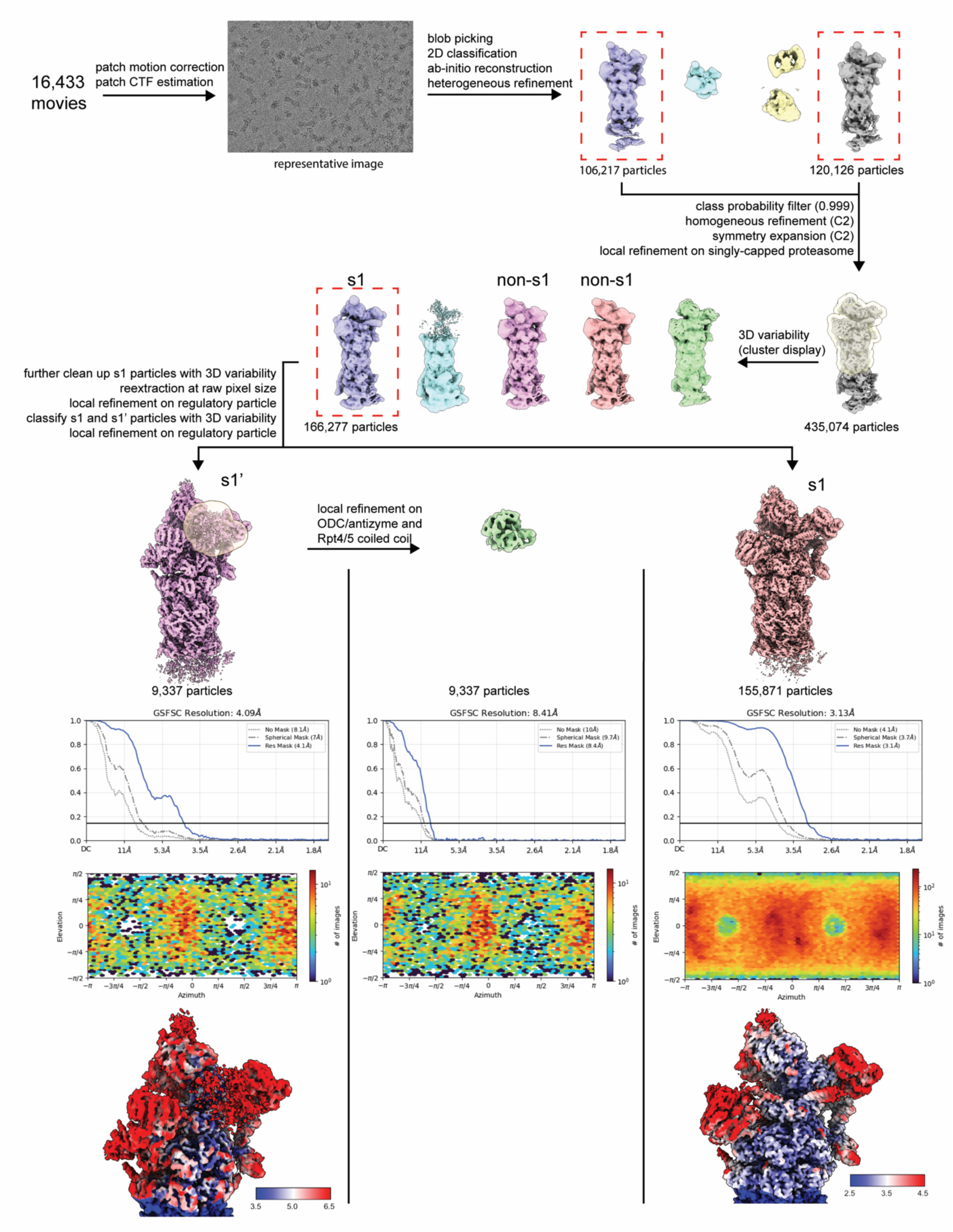
Cryo-EM data processing workflow for the proteasome bound to WT yODC/Antizyme. Schematic of the single-particle processing strategy used for the cryo-EM dataset collected on the 26S proteasome incubated with WT yODC/Antizyme. Particle counts are indicated at each stage, and final reconstructions are annotated with their conformational state. Gold-standard Fourier shell correlation curves (0.143 threshold), particle orientation distributions, and local-resolution estimates for each final EM volume are shown.

**Extended Data Fig 9.**
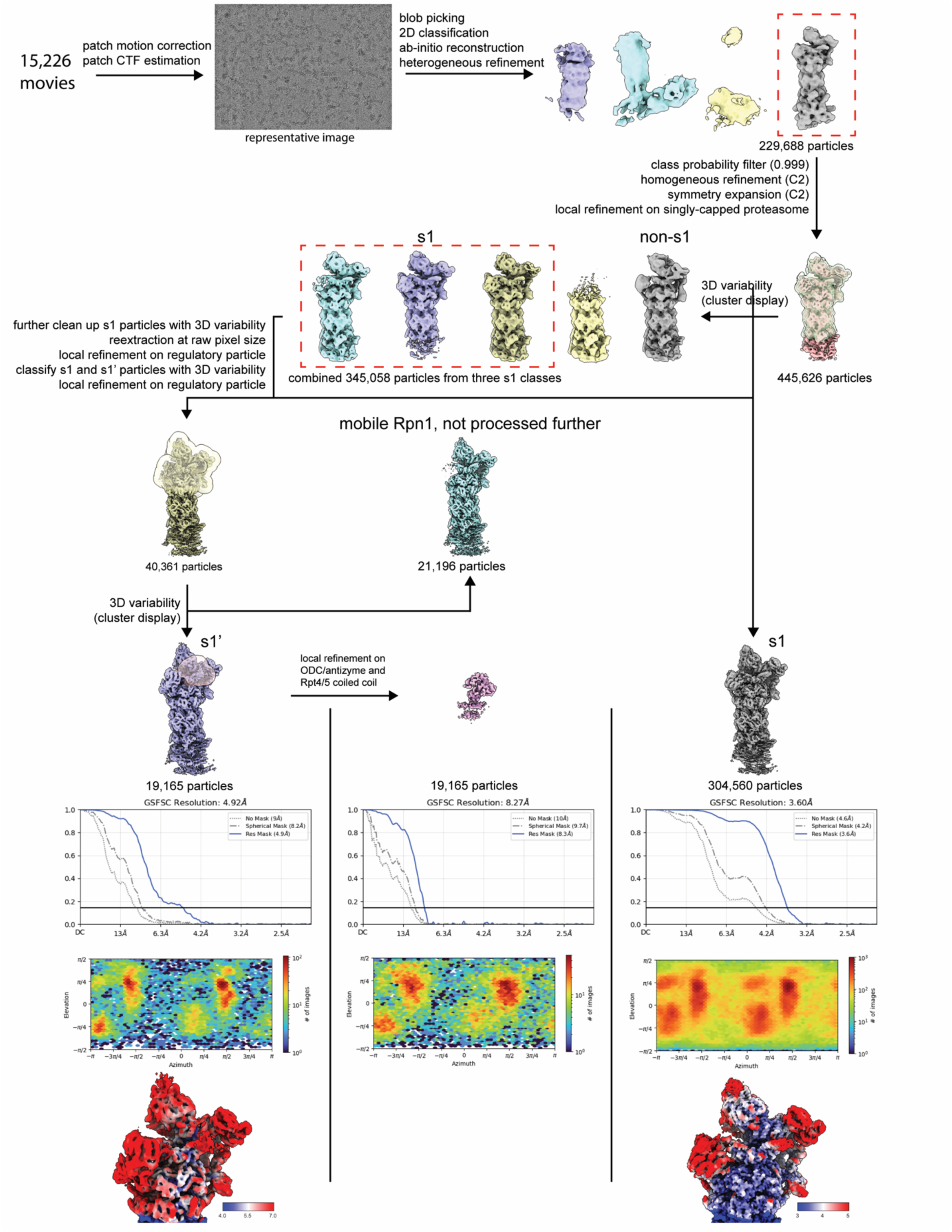
Cryo-EM data processing workflow for the proteasome bound to Δ34yODC/Antizyme. Schematic of the single-particle processing strategy used for the cryo-EM dataset collected on 26S proteasome incubated with Δ34yODC/Antizyme. Particle counts are indicated at each stage, and final reconstructions are annotated with their conformational state. Gold-standard Fourier shell correlation curves (0.143 threshold), particle orientation distributions, and local-resolution estimates for each final EM volume are shown.

**Extended Data Fig. 10.**
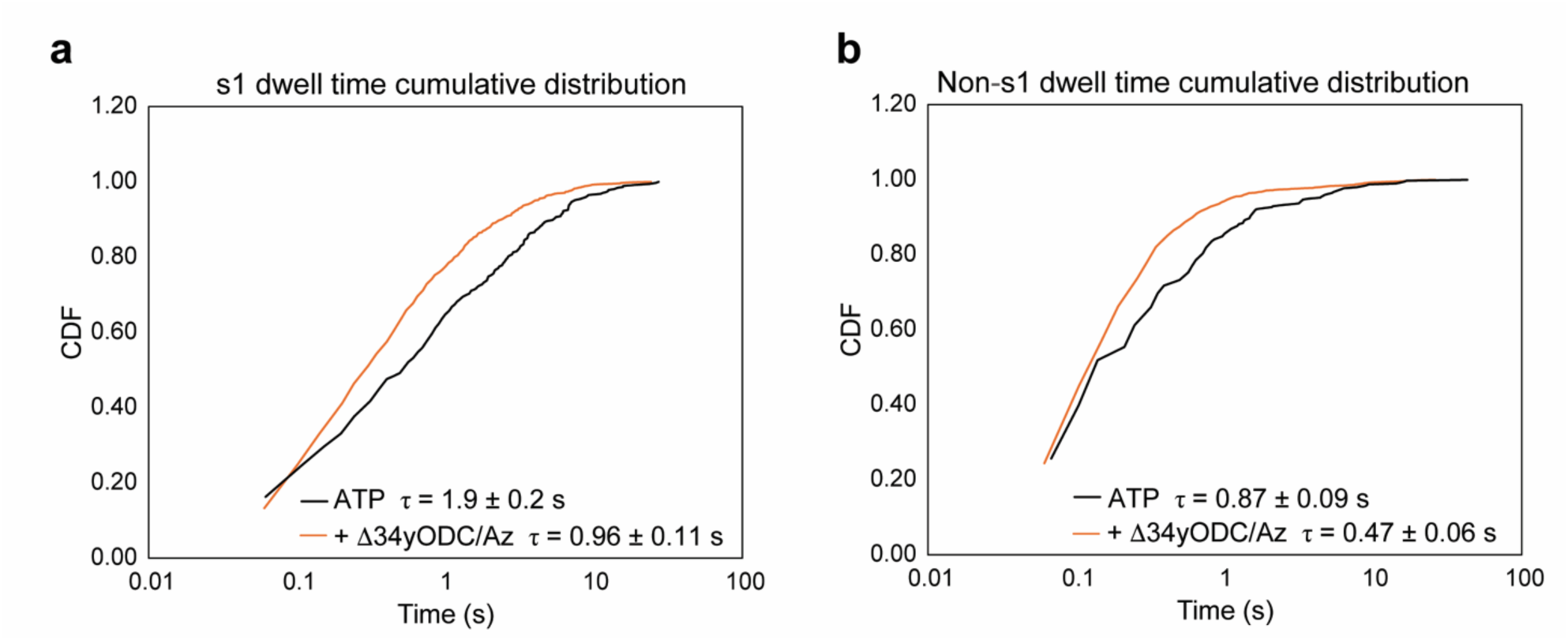
**yODC/Az binding affects the proteasome conformational dynamics. a-b**, Cumulative dwell time distributions of low-FRET s1/s1’ states (**a**) and high-FRET non-s1 states (**b**). The characteristic dwell times τ are reported as the mean of all observed dwell times. Number of single molecules used for analysis: 93 for ATP, 73 for Δ34yODC/Antizyme, 40 for Ub-substrate, and 34 for WT yODC/Antizyme.

